# Pup defence in lactating rats: The underlying neuropeptide signalling and their interactions in the nucleus accumbens shell

**DOI:** 10.1101/2024.10.29.620799

**Authors:** Alice Sanson, Annika Köck, Luisa Demarchi, Karl Ebner, Oliver J. Bosch

**Affiliations:** Department of Behavioural and Molecular Neurobiology, Regensburg Center of Neuroscience, University of Regensburg, Regensburg, Germany; Department of Pharmacology and Toxicology, Institute of Pharmacy and Center for Molecular Biosciences Innsbruck (CMBI), Leopold Franzens University Innsbruck, Innsbruck, Austria

## Abstract

Maternal aggression is a core feature of rodent maternal behaviour to defend their offspring from potential threats and is modulated by corticotropin-releasing factor (CRF) and oxytocin (OXT) systems signalling. Here, we investigated the involvement of those neuropeptide systems in maternal aggression within the nucleus accumbens shell (NAcSh), a central region of the reward and maternal circuits. Infusion of CRF or Urocortin3 (CRF-receptor 2 agonist), as well as an OXT receptor antagonist, reduced maternal aggression, suggesting a role in pup defence. Furthermore, the effects of CRF infusion in the NAcSh continued beyond the maternal defence test (MDT), reducing nursing and increasing self-grooming. Corroborating the involvement of the stress system in maternal aggression, colocalization of CRF and cFos immunoreactive cells were increased in response to the MDT, regardless of pup presence. In addition, MDT exposure increased intra-NAc OXT release in lactating rats, which could be also triggered by local retrodialysis of CRF, but not Urocortin3. However, both ligands of the CRF system elicited dopamine (DA) release in different dynamics. *Crh-r1 were* predominantly expressed in the medial NAc on medium-sized spiny neurons (MSN), but also in the rostral part on GABAergic interneurons. *Crh-r2* were mainly expressed in the rostral NAc and its expression on GABAergic interneurons increased towards the caudal pole. Lastly, we identified CRF-enriched projections to the NAcSh descending from the prefrontal cortex, the amygdala, and the paraventricular thalamus, among others.

In conclusion, intra-NAcSh dampened CRF system activity and enhanced OXT system transmission are indispensable for successful pup defence. Any perturbations like increased CRF system signalling might activate compensatory mechanisms to ensure adequate maternal behaviour.

## 1. Introduction

In numerous mammalian species, defence of their offspring from a potential threat, i.e., maternal aggression, is crucial to ensure the survival of the young (Svare, 1981). For the onset of maternal aggression in rodents, the behavioural response needs to shift from “flight” to “fight” (Gammie et al., 2008) and depends on factors such as the context, sensory inputs from the pups and the intruder, or the maternal hormonal profile (Bosch, 2013; de Almeida et al., 2014; Gammie and Lonstein, 2006; Lonstein and Gammie, 2002). Furthermore, decreased anxiety postpartum (Lonstein, 2007; Neumann, 2001) might be a prerequisite for enhanced aggressive behaviours towards a threat (Bosch, 2013; Lonstein and Gammie, 2002; Neumann and Bosch, 2008). However, fighting an intruder to protect the offspring elicits a stress response in the mother, and plasma ACTH and glucocorticoid levels increase during maternal aggression (Gammie et al., 2008; Neumann et al., 2001). Thus, the encounter with an unfamiliar intruder represents a psychosocial stressor for the dam (Bosch, 2013; Neumann et al., 2001). It is not surprising that neuropeptides related to stress and anxiety are implicated in maternal aggression (Bosch, 2013; Lonstein and Gammie, 2002), including the corticotropin-releasing factor (CRF) and the oxytocin (OXT) systems (Bosch, 2013; Gammie et al., 2008).

The CRF system comprises two receptor subtypes, CRF receptor (CRF-R) 1 and 2, and four ligands (CRF and Urocortins – UCN – 1, 2, 3), which bind to the CRF-Rs with different affinities (Deussing and Chen, 2018). Furthermore, the CRF family includes the CRF binding protein (CRF-BP), which mainly sequesters freely available CRF and UCN1 and prevents them from binding to CRF-R, thereby dampening their transmission (Ketchesin et al., 2017; Westphal and Seasholtz, 2006). The hypothalamic CRF-BP is a major factor for the reduced stress-responsiveness in lactating rats (Sanson et al., 2024c). CRF is primarily known to induce the physiological responses to stress by activating the hypothalamic-pituitary adrenal (HPA) axis (Deussing and Chen, 2018). However, both the CRF-R and their ligands are broadly expressed throughout the brain, and CRF-immunoreactive cells are present in several brain regions, including the nucleus accumbens (NAc) (Bangasser, 2013; Deussing and Chen, 2018; Eckenwiler et al., 2024). Hence, by binding to CRF-R within the brain, CRF exerts an anxiogenic and pro-depressive activity (Deussing and Chen, 2018). Furthermore, CRF-R signalling needs to be dampened during the peripartum period to ensure the full display of maternal behaviours (for review, see (Klampfl and Bosch, 2019a; Sanson et al., 2024a)), including maternal aggression (Gammie et al., 2008; Klampfl and Bosch, 2019a; Lonstein and Gammie, 2002; Sanson et al., 2024a). To this date, nothing is known about the role and interactions of the CRF and OXT systems in the NAc in the context of pup defence.

During the peripartum period, OXT signalling is crucial. This nonapeptide is mainly produced in magnocellular neurons of the paraventricular nucleus (PVN) and the supraoptic nucleus (SON) of the hypothalamus, which project to several regions of the social brain network (Menon and Neumann, 2023). OXT exerts well-known peripheral functions related to reproduction and, within the brain, it controls complex behaviours such as social and maternal behaviours via binding to its OXT receptor (OXT-R) (Jurek and Neumann, 2018; Menon and Neumann, 2023; Sanson and Bosch, 2022). In the peripartum period, OXT levels are increased to facilitate maternal responses (Pedersen, 1997; Rilling and Young, 2014; Yoshihara et al., 2018), and impaired OXT-R signalling can reduce maternal behaviour, including pup defence (for review, see (Sanson and Bosch, 2022; Sanson et al., 2024a)). Furthermore, OXT release in the PVN and central amygdala positively correlates with the mothers’ aggressive behaviour (Bosch et al., 2005).

The NAc belongs to the ventral striatum and is part of the brain reward circuit with dopaminergic (DA) inputs from the ventral tegmental area (VTA) (Floresco, 2015). This rather complex region can be distinguished into a central core (NAcCo) and an outer shell (NAcSh). The rat NAc has an additional level of organization as a neurochemical gradient that can be found along the rostro-caudal axis of this region (Mathieu et al., 1996; Ranaldi and Beninger, 1994; Rogard et al., 1993; Voorn and Docter, 1992), and DA receptor densities increase from the rostral to the caudal pole of the NAc (Allin et al., 1989). Besides some interneuron populations, appr. 95% of NAc cells are medium-sized spiny neurons (MSN), which express either DA receptor 1 or 2 and are mostly gamma-aminobutyric acid (GABA)ergic (Castro and Bruchas, 2019; Kardos et al., 2019; Robison and Nestler, 2011). Most NAc interneurons are GABAergic, expressing parvalbumin (Pvalb) or somatostatin and neuropeptide Y, whereas < 5% are cholinergic interneurons (Collins et al., 2019; Kardos et al., 2019). Furthermore, the NAcCo and NAcSh show differential connectivity, implying that they may control different behaviours (Brog et al., 1993; Floresco, 2015; Salgado and Kaplitt, 2015). Interestingly, in mice the NAc receives CRF inputs from stress-related regions, highlighting its involvement in stress-induced responses (Itoga et al., 2019). Beyond its role in the reward and stress circuits, the NAc regulates social behaviour via different neuropeptides and neurotransmitters (Borland, 2025) and is part of the complex maternal brain network, which modulates pup-directed responses and maternal behaviour (Dulac et al., 2014; Numan, 2007; Numan et al., 2005b; Servin-Barthet et al., 2023; Smiley et al., 2019; Stolzenberg et al., 2007). The NAcSh plays a specific role in maternal care, memory, and motivation (D’Cunha et al., 2011; Li and Fleming, 2003a, b; Numan et al., 2005a; Witchey et al., 2024) through OXT-R (Borland, 2025; D’Cunha et al., 2011; Witchey et al., 2024) and DA receptor D1 (Numan et al., 2005a), among others.

In this study, we aimed to unravel the involvement of the CRF and OXT systems in the NAcSh in the context of maternal aggression adopting a functional and a descriptive approach. Hence, we first tested if acute bilateral pharmacological modulations of these systems impair maternal aggression during the maternal defence test (MDT, a psychosocial stressor), as well as maternal care after exposure to this stressor. Using a descriptive approach, we characterized the molecular effects of MDT exposure at the level of the NAcSh by measuring mRNA and protein levels of CRF and OXT family members. As a next step, we investigated whether intra-NAc activation of CRF-R signalling affects local OXT and DA release. Lastly, we analysed the distribution as well as the nature of CRF-R-expressing cells in the NAcCo and NAcSh of lactating rats. In a separate cohort of virgin female rats, we further characterized the CRF system within the NAc by performing a retrograde tracing study to identify the major sources of CRF to the NAcSh in female rats.

## 2. Materials and methods

### 2.1. Animals

Virgin female Sprague-Dawley rats (230g - 250g at arrival; Charles River, Sulzfeld, Germany) were housed in groups of 3-4 rats of the same sex under standard laboratory conditions (12:12 h light/dark cycle; lights on at 07:00 am; room temperature 22 ± 2 °C, 55 ± 5% relative humidity), with *ad libitum* access to water and standard rat chow (ssniff-Spezialdiäten GmbH, Soest, Germany). To obtain lactating subjects, two virgin female rats were housed with a sexually experienced male Sprague-Dawley rat in Eurostandard type IV cages (40 × 60 × 20 cm) for 10 days. Afterwards, female rats were housed together in groups of 3-4 until potential pregnancy day 18 (PD18), when they were single housed in observational cages (38 × 22 × 35 cm) to ensure undisturbed delivery. On the day of birth (lactation day, LD0), litters were culled to 8 pups with balanced sexes. All rats from Experiments 1, 3 and 4 were handled daily to familiarize them with the experimenters, the procedures and to reduce non-specific stress responses.

Naïve virgin female Sprague-Dawley rats (200g - 220g at arrival; Charles River) at random stages of the estrous cycle were used as intruders for the MDT (Bosch, 2013). The intruders were housed in groups of 3-4 in a separate room to avoid olfactory recognition, which might influence the aggressive response of the resident dam (Klampfl et al., 2013). To visualize CRF projections to the NAcSh, virgin females were housed in groups of 2 until stereotaxic surgery and single housed in observational cages for 3 weeks thereafter.

The studies were conducted in accordance with the ARRIVE guidelines, the European regulations of animal experimentation (European Directive 2010/63/EU), and were approved by the local government of Unterfranken (Bavaria, Germany). According to the 3-Rs principles, all efforts were made to minimise the number of animals and their distress or suffering.

### 2.2. Experimental design

A graphical overview of the experimental design can be found in Figure 1A. All tests were performed during the light phase between 08:00 h and 13:00 h. Figure 1B shows a representative histological image of a correct implantation site (assessed as described in 2.6.).

**Figure 1.**
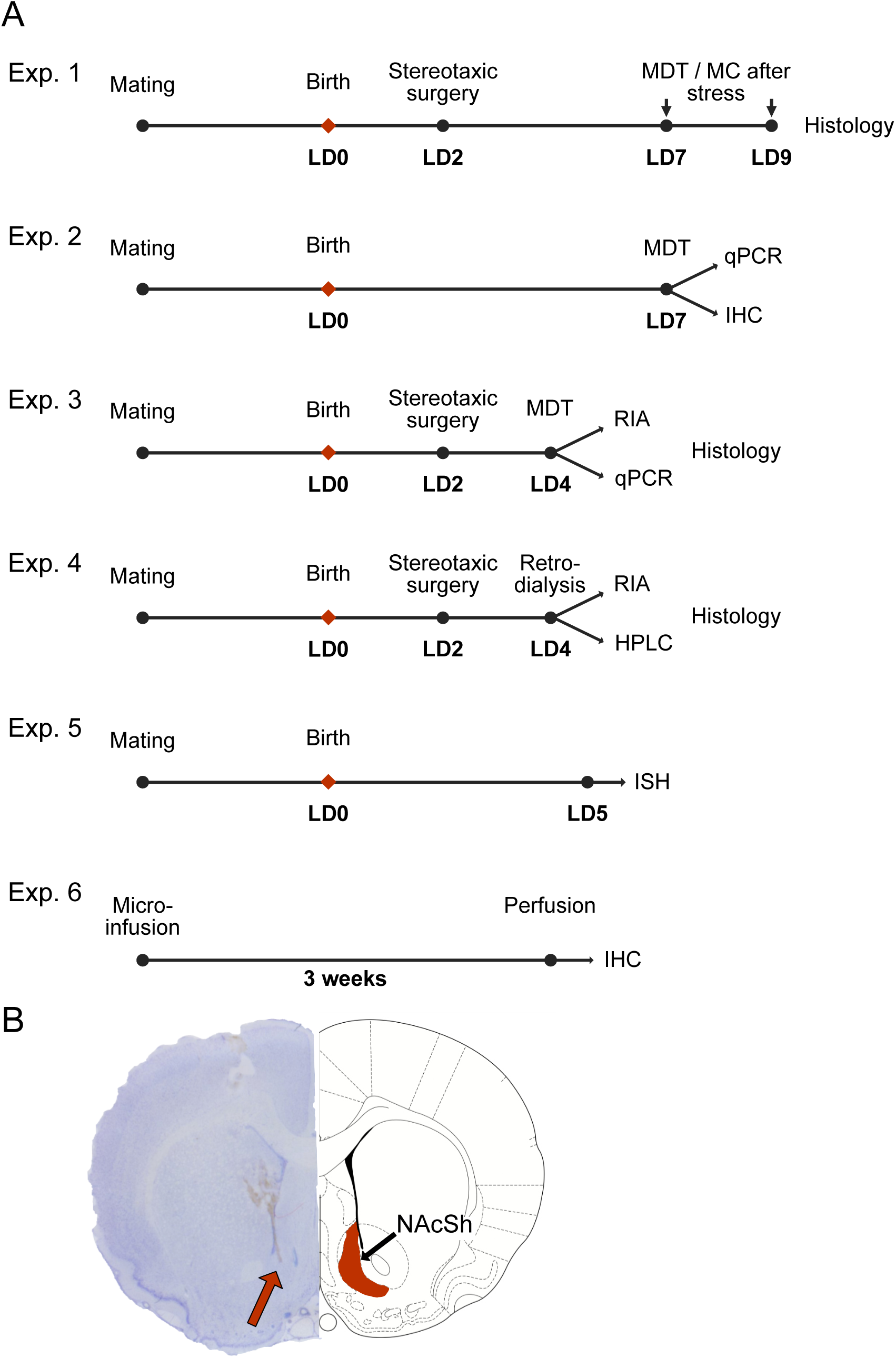
Experimental designs and representative histological picture of cannula/probe placement. (A) Experimental timelines for each experiment. (B) Representative histological picture of cannula/probe placement (left) and coronal scheme (right; (Paxinos and Watson, 2013)). Abbreviations: HPLC, high-performance liquid chromatography; IHC, immunohistochemistry; LD, lactation day; MC, maternal care; MDT, maternal defence test; qPCR, quantitative real-time polymerase chain reaction; RIA, radioimmunoassay.

#### 2.2.1. Functional approach: behavioural observations following NAcSh neuropeptides’ manipulation

##### Experiment 1

Maternal aggression (see 2.3.2.) was tested in the home cage for 10-min during the MDT following pharmacological manipulations of the CRF and OXT family members, i.e., by acute local, bilateral infusion of agonists and antagonists of the CRF-R1, - R2, inhibitor of CRF-BP, or OXT-R antagonist (Table 1) via previously implanted guide cannulas (see 2.4.1.). Doses, lag time and corresponding VEH were chosen according to previous studies (Bosch et al., 2005; D’Anna and Gammie, 2009; Klampfl et al., 2016; Klampfl et al., 2018; Sanson et al., 2024c). As CRF binds to CRF-R1 with 40x higher affinity than to CRF-R2 (Deussing and Chen, 2018), it is considered to mainly act on this subtype and the same dose was previously used as a selective agonist of CRF-R1 (Klampfl et al., 2018). However, we tested for CRF potentially acting on CRF-R2 by pre-treatment with a selective CRF-R1 antagonist followed by an injection of either vehicle (VEH) or CRF 10 min after the first infusion. To reduce the number of animals used, the same rat underwent the MDT on LD7 and on LD9 after treatment with different drugs (Figure 1A) and, to control for any long-lasting effects of the LD7 treatments, the treatments of LD9 were balanced and mixed. In addition, undisturbed maternal care was observed for 60 min preceding the MDT and pharmacological manipulations, as well as following the combined MDT and drug treatment. At the end of the MDT, mothers were anesthetised with CO_2_ and subsequently decapitated. Their brains were extracted, flash-frozen in n-methylbutane, and stored at −80°C until estimation of correct implantation as described in 2.6.

**Table 1.**
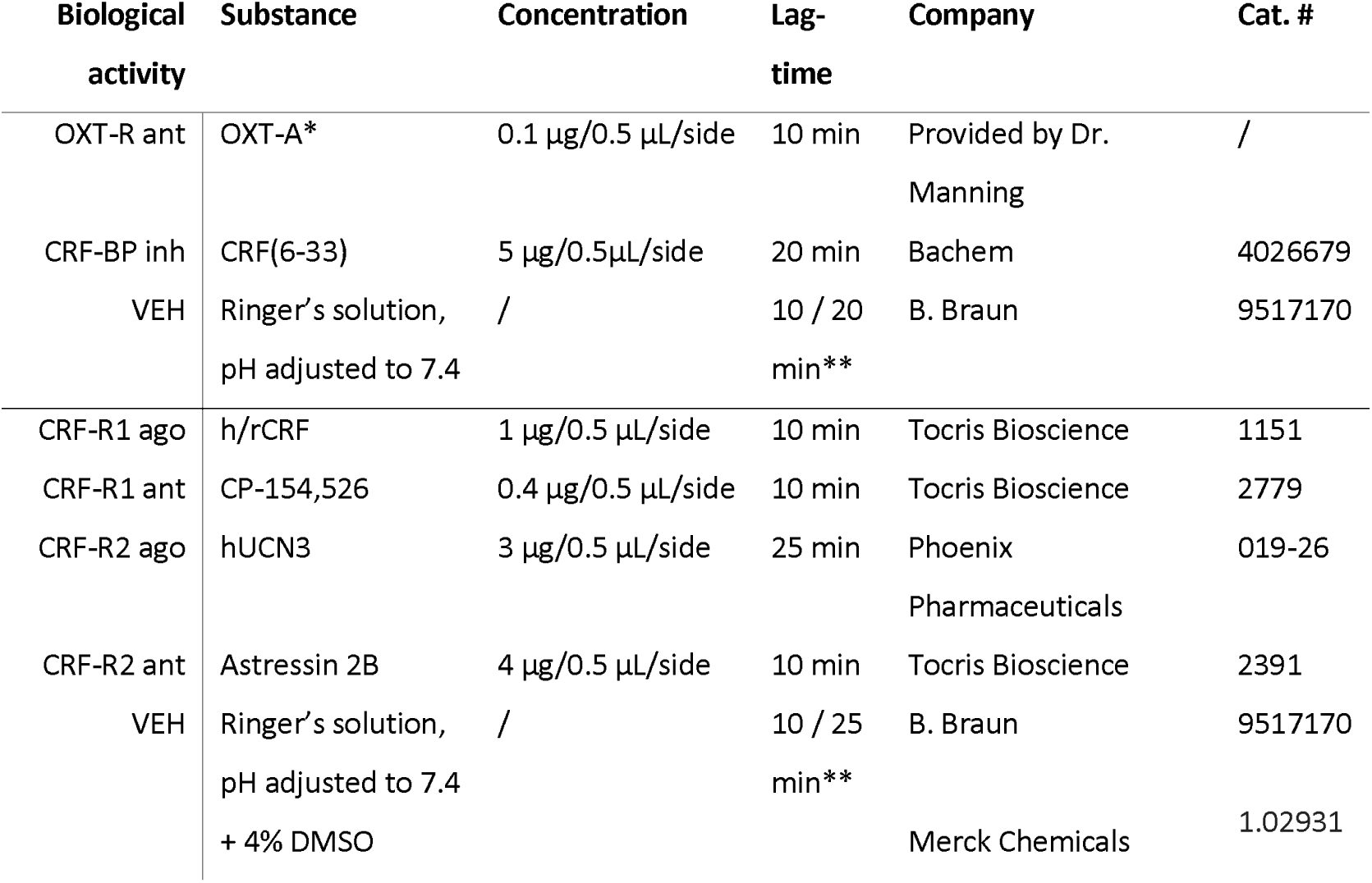
Details of administered drugs. Abbreviations: ago, agonist; ant, antagonist; inh, inhibitor; VEH, vehicle. * OXT-A: ((d(CH_2_)_5_^1^, Tyr(Me)^2^, Thr^4^, Orn^8^, des-Gly-NH_2_ ^9^)-vasotocin) ** Depending on the corresponding infused substance.

| Biological activity | Substance | Concentration | Lag-time | Company | Cat. # |
| --- | --- | --- | --- | --- | --- |
| OXT-R ant | OXT-A* | 0.1 µg/0.5 µL/side | 10 min | Provided by Dr. Manning | / |
| CRF-BP inh | CRF(6-33) | 5 µg/0.5µL/side | 20 min | Bachem | 4026679 |
| VEH | Ringer's solution, pH adjusted to 7.4 | / | 10 / 20 min** | B. Braun | 9517170 |
| CRF-R1 ago | h/rCRF | 1 µg/0.5 µL/side | 10 min | Tocris Bioscience | 1151 |
| CRF-R1 ant | CP-154,526 | 0.4 µg/0.5 µL/side | 10 min | Tocris Bioscience | 2779 |
| CRF-R2 ago | hUCN3 | 3 µg/0.5 µL/side | 25 min | Phoenix Pharmaceuticals | 019-26 |
| CRF-R2 ant | Astressin 2B | 4 µg/0.5 µL/side | 10 min | Tocris Bioscience | 2391 |
| VEH | Ringer's solution, pH adjusted to 7.4 | / | 10 / 25 min** | B. Braun | 9517170 |
|  | + 4% DMSO |  |  | Merck Chemicals | 1.02931 |

#### 2.2.2. Descriptive approach: neurobiological characterization of the NAcSh

##### Experiment 2

MDT was performed on LD7 ± 1 in the absence of surgical procedures (Figure 1A) to analyse the central changes of the CRF and OXT systems (see 2.7.). Rats were either exposed to the MDT or not (“No MDT”). To assess potentially stress-dampening effects of the pups’ presence, these groups were subdivided into pups being present in the home cage throughout the test (“with pups”) or removed immediately after the MDT (“without pups”) resulting in the following 4 groups: 1. *No MDT - with pup s*(control group); 2. *MDT - with pups*; 3. *No MDT - without pups*; 4. *MDT - without pups*.

Maternal care (2.3.1.) was assessed in all groups for 60 min under undisturbed conditions prior to MDT testing, which was performed as described in 2.3.2., followed by further 60 min of maternal care evaluation after stress exposure (*MDT - with pups*) and in the control group *No MDT - with pups*. 90 min after the undisturbed maternal care assessment (i.e., 80 min after termination of the MDT), all dams were deeply flash anesthetised with Isoflurane (Baxter Germany GmbH, Unterschleißheim, Germany) before cardiac perfusion with ice-cold 1X phosphate-buffered saline (PBS) and were subsequently decapitated. Their brains were then flash-frozen in n-methylbutane and stored at −80°C until bisecting into the two hemispheres for further analysis (see 2.7.). Left and right hemisphere were randomly assigned to qPCR or immunofluorescence analysis.

##### Experiment 3

Microdialysis was performed on LD4 ± 1 to measure OXT release during and after exposure to the MDT (see 2.3.2.). 100 min after the end of the MDT, mothers were anesthetised with CO_2_ and subsequently decapitated. Their brains were extracted, flash-frozen in n-methylbutane, and stored at −80°C until further processing. The correct implantation of microdialysis probes was assessed as described in 2.6. To further characterize the physiological changes in OXT levels during the MDT, punches of the PVN and SON were harvested to potentially correlate OXT mRNA levels to aggressive behaviour as well as OXT release within the NAc (2.7.2.).

##### Experiment 4

To measure OXT or DA release in the NAc in response to continuous local CRF-R activation, retrodialysis was performed on LD4 ± 1 under non-stress conditions (Figure 1, see also 2.4.). For OXT or DA measurements, separate cohorts of rats were used.

##### Experiment 5

To characterize the distribution and expression pattern of *Crh-r1* and *- r2* along the rostro-caudal axis of the NAc during lactation, local mRNA expression was visualized via fluorescent *in-situ* hybridization (ISH) (RNAscope^®^, Advanced Cell Diagnostics Inc., CA, USA). On LD 5 ± 1, lactating female rats were deeply anaesthetised before they underwent cardiac perfusion as described for Experiment 2, followed by perfusion with 4% PFA in 1X PBS. Brains were collected and post-fixed in the same fixative, overnight at 4°C. Afterwards, the brains were transferred to a 30% sucrose solution for cryoprotection and stored at 4°C until they sank. Finally, the brains were flash-frozen in ≥ 99% n-methylbutane and stored at −80°C until ISH was performed (see 2.7.3.).

##### Experiment 6

To visualize the CRF afferents projecting to the NAcSh, a retrograde transported virus was unilaterally microinfused in the right NAcSh of virgin female rats (n = 2; see 2.5). After stereotaxic microinfusion, rats were left undisturbed for 3 weeks to allow viral retrograde transport and expression. Then, rats were deeply anesthetised and underwent cardiac perfusion as described for Experiment 5. Finally, brains were collected and flash-frozen in n-methylbutane and stored at −20°C until further processing (as described in 2.7.4.).

### 2.3. Behavioural assessment

#### 2.3.1. Maternal care

After 60 min of habituation to the experimental room, maternal care (MC) was monitored under undisturbed conditions as well as after drug infusion (Experiment 1) under stress conditions, *i.e.*, following the MDT. According to an established protocol, an experimenter blind to the treatment observed the behaviour for 10s every second minute, in 30 min blocks (Bosch and Neumann, 2008). Different behaviours related to pup care were quantified. The main parameters indicating the quality of maternal behaviour were arched-back nursing (ABN) and licking and grooming (LG) (Bosch, 2011; Klampfl and Bosch, 2019b). Other nursing parameters included passive nursing postures such as blanket posture, nursing laying on the side or back, and hovering over the pups, which together account for “total nursing”. Along with maternal behaviours, non-maternal behaviours were quantified (off nest), including self-grooming.

#### 2.3.2. Maternal aggression

The MDT was performed in the presence of the litter using a modified resident-intruder test (Klampfl et al., 2018). A virgin female intruder was placed into the home cage of the mother (resident) for 10 min (Klampfl et al., 2018). In Experiment 1, the MDT was performed after drug infusion and the corresponding lag time (see Table 1). The behaviour of the mother was recorded and analysed with JWatcher (http://www.jwatcher.ucla.edu/) by an experimenter blind to the treatment. The following parameters for aggressive behaviour were scored: number of attacks, latency to first attack, number and time spent in lateral threat, pinning, offensive upright, aggressive sniffing and grooming (Bosch, 2013). Moreover, non-aggressive behaviours were monitored, including intruder exploration, self-grooming and resting. To provide a more precise behavioural phenotyping, z-score normalization for maternal aggression was performed (for details, see section 2.8.). The behavioural ethograms during the MDT were generated with Boris v.8.27.1 (Friard and Gamba, 2016).

### 2.4. Local infusions, micro- and retrodialysis

#### 2.4.1. Stereotactic surgeries

Rats underwent surgical procedures on LD2 ± 1. Stereotaxic surgeries were performed under semi-sterile conditions under inhalation anaesthesia as previously described (Bosch et al., 2010). Briefly, for local infusions, 23 G guide cannulas (length: 12 mm) were implanted bilaterally 2 mm above the NAcSh (coordinates: anterior posterior +1 mm, lateral ± 3 mm, ventral - 5.3 from the skull surface, angle 17.5° (Paxinos and Watson, 2013)).

Probes for micro- and retrodialysis were identical, as was their implantation (Bosch et al., 2004; Klampfl et al., 2018). Briefly, probes (molecular cut-off: 18 kDa; Hemophan, Gambro Dialysatoren, Hechingen, Germany) were implanted unilaterally targeting the right NAc on LD2 ± 1, adopting the surgical procedure described above (coordinates: anterior posterior +1 mm, lateral ± 3 mm, ventral - 7.3 from the skull surface, angle 17.5° (Paxinos and Watson, 2013)). Due to the size of the micro- / retrodialysis probes, a distinction between NAcSh and NAcCo was not feasible, thus we targeted the whole NAc.

#### 2.4.2. Local infusion

For acute local infusion, 30 G infusion cannulas (length: 14mm) were connected via a PE-50 tubing to 10 µL Hamilton syringes. The infusion cannula was lowered in the guide cannula and kept in place for approximately 30 s during drug infusion. The substances were bilaterally infused in a volume of 0.5 µL / side and are referenced in Table 1.

#### 2.4.3. Microdialysis procedures

Microdialysis was performed as previously described (Bosch et al., 2004; Klampfl et al., 2018). Briefly, on the test day, the inflow adapter of the microdialysis probe was connected via PE-20 tubing to a syringe mounted onto a microinfusion pump, while the outflow adapter was attached to a 1.5 mL collection tube. The probe was flushed at a rate of 3.3 µL/min with sterile Ringer’s solution (pH adjusted to 7.4) for 90 min before the start of sample collection. Starting at 10 a.m., samples 1 and 2 were collected under basal conditions (Ringer’s solution infusion) in 30-min intervals. Sample 3 was collected during exposure to a virgin female intruder for 10 min (see 2.3.2.) and an additional 20 min thereafter, followed by collection of samples 4 and 5 in 30-min intervals. All samples were immediately frozen on dry ice and stored at −20°C until further measurements (2.4.5.).

#### 2.4.4. Retrodialysis procedures

On the test day, rats were connected to the syringes as described in 2.4.3. The probe was flushed at a rate of 1 µL/min with sterile Ringer’s solution (pH adjusted to 7.4) for 90 min before the start of sample collection as previously described (Klampfl et al., 2018). For OXT measurement, starting at 10 a.m., samples 1 and 2 were collected under basal conditions (infusion of Ringer’s solution, pH adjusted to 7.4) in 30-min intervals. Afterwards, the syringes containing Ringer’s solution were exchanged with those filled with either CRF (0.05 µg/µL) or UCN3 (0.15 µg/µL) and drugs were infused for 30 min (sample 3; doses based on (Klampfl et al., 2018)) at a continued rate of 1 µL/min. Syringes were then switched back to those containing sterile Ringer’s solution and samples 4 and 5 were collected in 30-min intervals. All samples were immediately frozen on dry ice and stored at −20°C until further measurements (2.4.5).

For DA measurement, all samples were collected in 10-min intervals in 1.5 mL tubes containing a stabilizing solution (sterile Antioxidant-Ascorbic acid solution; based on (Ebner et al., 2022), with minor adaptations). Starting at 10 a.m., samples 1-8 were collected under basal conditions (infusion of Ringer’s solution, pH adjusted to 7.4). Afterwards, the syringes were exchanged with those filled with the drugs as described above, which were infused for 30 min (samples 9-11). Syringes were then switched back to those containing sterile Ringer’s solution, and samples 12-15 were collected. All dialysate samples were immediately frozen on dry ice and stored at −80°C until further measurements (see 2.4.6).

#### 2.4.5. Radioimmunoassay

OXT content in the samples of Experiments 3 and 4 was measured by an external company following lyophilisation using a highly sensitive and selective radioimmunoassay (RIA, detection limit: 0.1 pg/sample; cross-reactivity of the antisera with other related peptides < 7%; RIAgnosis, Sinzing, Germany). OXT content was expressed as pg in 100 µL dialysate (microdialysis) or as pg in 30 µL dialysate (retrodialysis) and reported as % change from baseline.

#### 2.4.6. High-performance liquid chromatography (HPLC)

DA content was measured by reverse-phase high-performance liquid chromatography (RP-HPLC) and the results were analysed as previously described ((Camats-Perna et al., 2019; Ebner et al., 2022); detection limit: 0.25 fmol/5 µL sample). The DA content in each dialysate was expressed as a relative value to the mean absolute DA content of the eight basal samples.

### 2.5. Microinfusion

70 nL of a retrograde transported virus (AAVrg-hSyn1-mCherry-WPRE, Vector Biolabs, Malvern, USA) was unilaterally microinfused in the right NAcSh (coordinates: anterior posterior + 1.7, lateral + 0.9, ventral - 6.4 from the brain surface (Paxinos and Watson, 2013) of virgin female rats (n = 2). To visualize the injection site, a GFP-associated reporter (AAV9-hSyn-eGFP-WPRE, Vector Biolabs) was mixed 1:1 with the AAVrg virus.

### 2.6. Histology

At the end of all pharmacological and micro-/retrodialysis experiments, rats were killed by an overdose of CO_2_ inhalation. Brains were sectioned in 40 µm coronal sections using a cryostat (CM3050S; Leica Microsystem GmbH, Nussloch, Germany), mounted on slides and Nissl stained to visualise the brain structures and to identify the implantation sites. Only animals with a correct placement of the cannulas / microdialysis probes were included in the statistical analysis.

### 2.7. Molecular analysis

#### 2.7.1. Neurobiological changes following the MDT: Immunofluorescence on slides

Brains from Experiment 2 were divided into their hemispheres, and one hemisphere was cut into coronal sections of 16 μm using a cryostat. Per rat, 7 sections containing the NAcSh were collected on SUPERFROST^®^ microscope slides (VWR, Radnor, USA) and stored at −20°C until further processing according to an established protocol (Demarchi et al., 2024). The slides were incubated overnight at 4°C with primary antibodies (rabbit anti-cFos 1:2000, #AB190289, Abcam; goat anti-CRF 1:1000, #Sc-1761, Santa Cruz) diluted in blocking solution (3% Bovine serum albumin, Sigma-Aldrich, Burlington, USA; 0.3% Triton X-100, Sigma-Aldrich; in PBS 1X). The next day, slides were washed in 1X PBS (3 × 10 min washing steps) and went through 2 h incubation at room temperature with secondary antibodies diluted in blocking solution (Alexa Fluor 488-conjugated anti-rabbit IgG 1:800; Alexa Fluor 594-conjugated anti-goat IgG 1:500). After 3 × 10 min washing steps in PBS 1X, the slides were finally mounted with ROTI^®^Mount FluorCare DAPI (Carl Roth Gmbh & Co. KG, Karlsruhe, Germany). For image analysis, all stained images of the NAcSh were acquired with 0.5 µm z-step size using a Thunder Leica microscope (Leica Thunder DM6 B, Camera Leica DFC9000 GT). To cover most of the NAcSh surface and to have an overview of the whole region, a total of 6 tiles per NAcSh were taken at 20X magnification and merged. In total, 7 images of the whole NAcSh were acquired per rat. Images were analysed manually using Fiji for ImageJ (Schneider et al., 2012) by an experimenter blind to the treatment conditions. Specifically, an average intensity projection was created, and the contrast was adjusted at 0.1% for each channel of each picture. The nuclei of the cells marked with DAPI were counted automatically. Only the cells marked with c-Fos and/or CRF that colocalized with DAPI were counted. The percentage of c-Fos and CRF co-labelled cells was calculated. The brightness of the representative images was adjusted using Fiji for ImageJ.

#### 2.7.2. Neurobiological changes following the MDT: Real-time qPCR

The other hemispheres from Experiment 2 as well as brains from Experiment 3 were cut into coronal sections of 250 µm using a cryostat. The whole NAc (Experiment 2; 1.70 – 0.70 from bregma (Paxinos and Watson, 2013); technical limitations did not allow us to select the NAcSh only), or the PVN and SON (Experiment 3; −1.08 – −1.80 from bregma (Paxinos and Watson, 2013)) were harvested using a 1-mm-diameter puncher and stored at −80°C until further processed. Total RNA was isolated using peqGold Trifast (VWR International, Radnor, United States) according to the manufacturer’s protocol. For mRNA analysis, 250 ng of total RNA per sample was reverse transcribed using Ultra Script 2.0 (PCR Biosystems, London, United Kingdom). Relative quantification of RNA levels was performed using PowerUp SYBR Green Master Mix (Thermo Fischer, Waltham, USA), and Glyceraldehyde-3-phosphate-dehydrogenase *(Gapdh)* was used as housekeeping gene. The targets and the sequences of all genes are listed in Table 2. Data were analysed following the 2^ΔCt method.

**Table 2.**
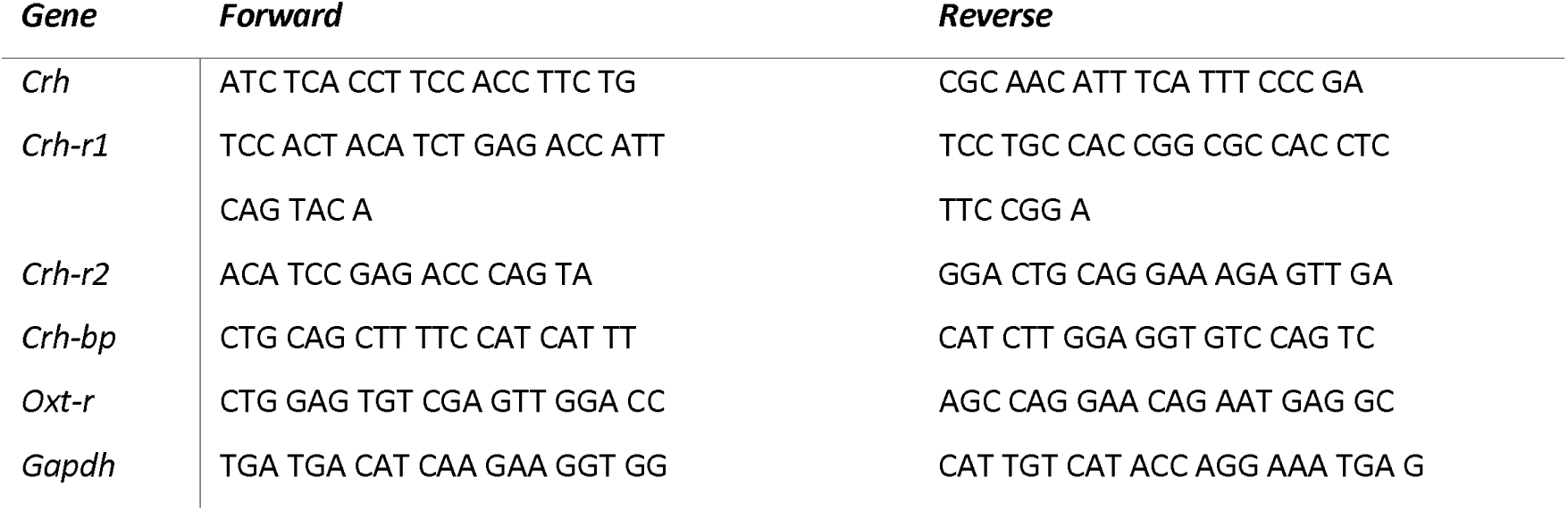
Primers forward and reverse sequences.

#### 2.7.3. Characterization of CRF-R-expressing cells: Fluorescent *in-situ* hybridization

Brains from Experiment 5 were cut into coronal slices of 16 μm using a cryostat (from bregma +1.70 to +0.70, (Paxinos and Watson, 2013)) and subsequently mounted on SUPERFROST^®^ Plus microscope slides (VWR). 2 slices per rat (4 x NAc) from the rostral, central, and caudal plane of the rat NAc were processed in a multiplex fluorescent v2 assay according to the manufacturer’s instructions (as detailed in #UM323100, Advanced Cell Diagnostics Inc., Newark, USA). The following probes were purchased from Advanced Cell Diagnostics Inc.: Rn-Crh-r1-C1 (Cat. #: 318911), Rn-Crh-r2-C1 (Cat. #: 417851), Rn-Ppp1r1b-C2 (Cat. #: 1048941-C2), Rn-Pvalb-C3 (Cat. #: 407821-C3).

4 × 5 tile images covering the left and right NAcCo and NAcSh were acquired at 20X magnification on a Leica Thunder DM6 B microscope using a Leica DFC9000 GT camera. This resulted in a total of 4 images of the whole NAc per plane and rat. Image analysis was semi-automatically performed using QuPath (version 0.5.1, (Bankhead et al., 2017)). For each channel, single measurement classifiers were created based on the integrated cell detection feature of the software. In order to distinguish between NAcCo and NAcSh, the region of interest was annotated in the software according to landmarks from the rat brain atlas (Paxinos and Watson, 2013). As a first step, we determined the overall *Crh-r* distribution along the rostro-caudal axis of the NAc by calculating the % of total *Crh-r*-expressing cells on the total count of DAPI cells by plane. For each NAc plane and subregion, we assessed the neurochemical properties of cells expressing *Crh-r* by calculating the % of *Crh-r* colocalization with DAPI and other markers on the total count of cells expressing *Crh-r* and DAPI regardless of co-expression with other markers (*Pvalb* and *Ppp1r1b*). We then calculated the average of all 4 NAc for each rat and plane and based on this, we determined the average including all 3 rats. Finally, we computed the % of NAc cells expressing a specific marker in each plane by calculating the average of each plane divided by the total count for the whole NAc. For all ISH experiments, ‘n’ represents the number of animals. The brightness and contrast of the representative images was adjusted using Fiji for ImageJ.

#### 2.7.4. Identification of CRF afferents: Free-floating immunofluorescence

To identify the main source of CRF, coronal slices of 40 µm were immunolabeled for CRF and mCherry. For each rat, five series were collected (from bregma +5.64 to −7.08, (Paxinos and Watson, 2013)) in a 24-well plate containing anti-freezing solution. Next, one series was processed for free-floating immunofluorescence to visualize mCherry and CRF using an established protocol (Menon et al., 2018). After blocking, slices were incubated overnight at 4°C with primary antibodies (chicken anti-mCherry 1:1000, #Ab205402, Abcam; goat anti-CRF 1:1000,, #Sc-1761, Santa Cruz) diluted in blocking solution (3% Bovine serum albumin; 0.3% Triton X-100; in PBS 1X). The following day, slices were washed 3 × 10 min in 1X PBS and went through 2 h incubation at room temperature with secondary antibodies (Alexa Fluor 594-conjugated anti-chicken IgG 1:1000; Alexa Fluor 488-conjugated anti-goat IgG 1:500) diluted in blocking solution. Then, slices underwent 3 × 10 min washing steps in PBS 1X + 0.3% Triton X 100, followed by 1 × 15 min washing in PBS 1X. The slides were finally mounted with ROTI^®^Mount FluorCare DAPI, and images of all sections were acquired using Leica Thunder DM6 B microscope with Leica DFC9000 GT camera at 10X and 20X magnification. The contrast was adjusted at 0.1% for each channel of each picture and brightness was adjusted for the representative images.

### 2.8. Statistical analysis

Statistical analysis was conducted using GraphPad Prism10 (GraphPad Software, Boston, USA). Data were tested for normality using Shapiro-Wilk test, homogeneity of variance was tested with F-test, and statistical outliers were identified with the Grubbs’ method and excluded from subsequent analysis. Where normality was violated, non-parametric tests were used; if homogeneity of variance was violated, we used the corresponding corrections. Data of maternal aggression and OXT release in response to the MDT were analysed using unpaired t-test, or non-parametric Mann-Whitney U test, or unpaired t-test with Welch’s correction. The temporal pattern of OXT release in response to the MDT was analysed with 1-way repeated measures ANOVA (factor: time); maternal care and retrodialysis data were analysed using 2-way ANOVA or Mixed-models analysis for repeated measures (factors: time and treatment); qPCR and immunofluorescence data were analysed using 2-way ANOVA (factors: pups and test). Where appropriate, post-hoc comparisons were performed using Bonferroni’s correction. Correlations between maternal aggression and maternal care or molecular data were performed using Pearson’s or Spearman’s correlations. Size effects were calculated with Cohen’s *d* coefficient and eta squared η^2^. Normalization of data using the z-score method was performed for maternal aggression, providing a more precise behavioural phenotyping (Begni et al., 2020; Guilloux et al., 2011). In detail, z-score for maternal aggression was calculated as follows (x: individual value; µ: mean of control group; σ: standard deviation of control group), as we previously described (Sanson et al., 2024c):

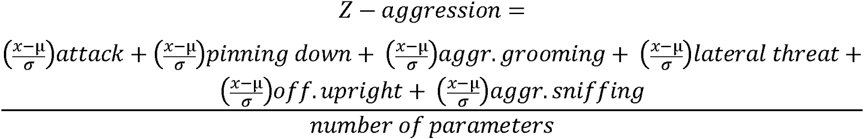

In each case, p ≤ 0.05 was considered statistically significant and a trend was accepted up to p = 0.07.

## 3. Results

### 3.1. Functional approach: Altered CRF and OXT transmission in the NAcSh impaired aspects of maternal behaviour under stress conditions

#### 3.1.1. CRF-R1 activation impaired maternal aggression and care under stress conditions

The experimental timeline is shown in Figure 2A. The representative ethograms (Figure 2B) of one VEH- and one CRF-treated mother indicate that mostly non-aggressive behaviours were displayed after CRF infusion. Indeed, CRF induced a significant reduction in the number of attacks (U = 48, p < 0.05; Mann Whitney test; Cohen’s *d* = 0.97; Figure 2C), but the latency to first attack (Figure 2D) and the overall z-score for maternal aggression (Figure 2E) were not altered (p > 0.05, each). Interestingly, also the time spent exploring the intruder was not altered by CRF treatment (p > 0.05; Figure 2F), while the time spent self-grooming during the MDT was significantly higher compared to VEH-treated dams (t[18]= 2, p < 0.05; Cohen’s *d* = 0.91; Table 3).

**Figure 2.**
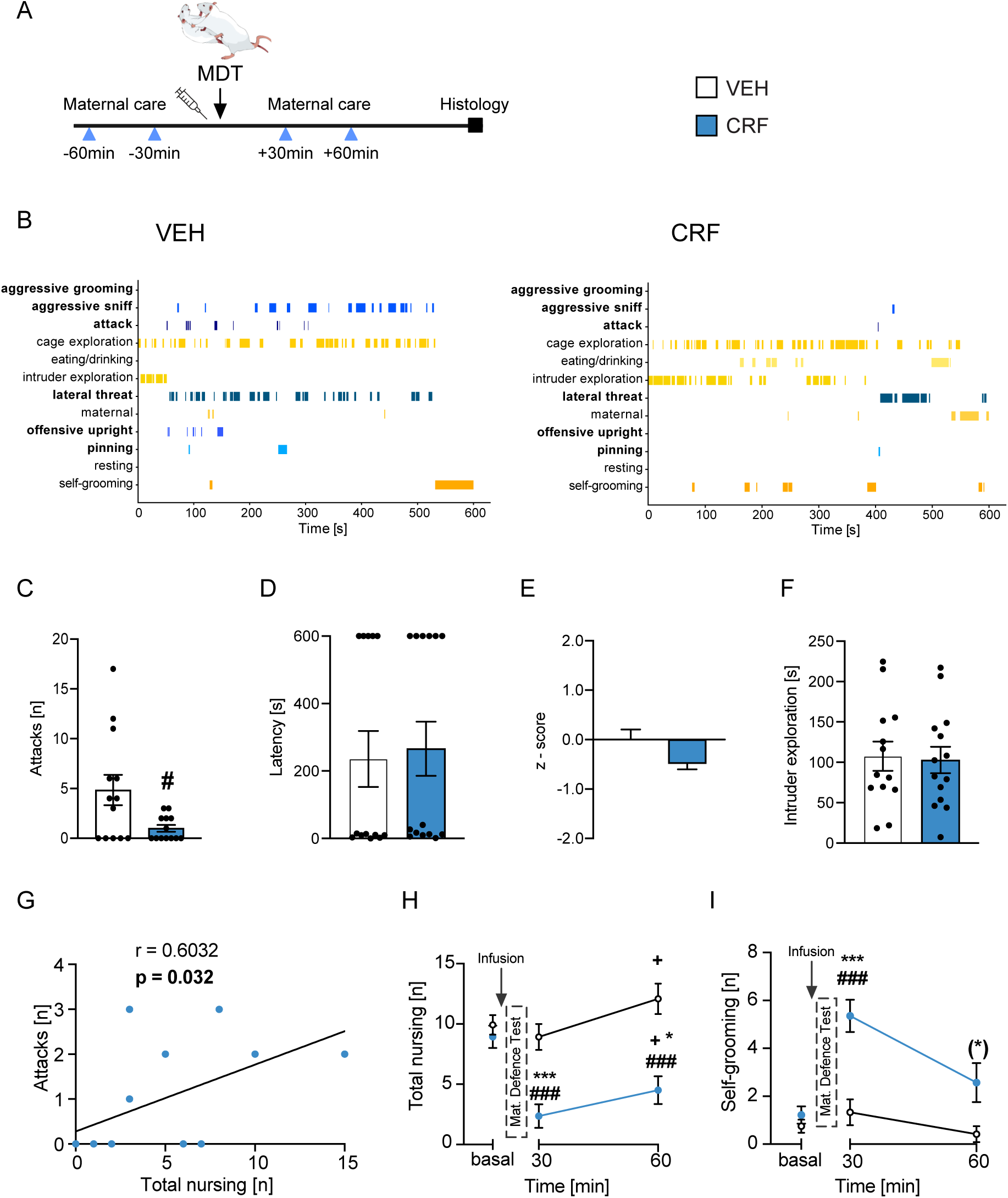
Infusion of CRF in the NAcSh impaired maternal aggression and care under stress conditions in lactating rats. (A) Experimental timeline (for details, see main text 2.2.1.). (B) Representative behavioural ethograms during the MDT of one dam, each, treated with vehicle (VEH) or CRF; aggressive behaviours are highlighted in bold and represented by blue bars. Behavioural parameters measured during the MDT: (C) number of attacks, (D) latency to first attack, (E) overall z-score for aggression, (F) time spent exploring the intruder; # p < 0.05 vs VEH-treated dams, Mann-Whitney test. (G) Spearman’s correlation between the number of attacks and total nursing at t60. Maternal care before (basal) and after treatment infusion combined with exposure to the MDT shown as (H) total nursing, and (I) self-grooming. ### p < 0.001 vs VEH-treated dams; (*) p ≤ 0.07, * p ≤ 0.05, *** p < 0.001 vs respective basal; + p < 0.05 vs respective previous time point; 2-way repeated measures ANOVA followed by Bonferroni post hoc comparisons.

**Table 3.** Effects of pharmacological manipulations within the NAcSh on aggressive and non-aggressive behaviours during the MDT in lactating rats. The occurrence and total time spent on aggressive and non-aggressive behaviour are expressed as mean ± SEM ((#) p ≤ 0.07, # p ≤ 0.05, ## p < 0.01 vs Vehicle (VEH)-treated dams; Unpaired t-test, or non-parametric Mann-Whitney U test, or unpaired t-test with Welch’s correction. The occurrence of attacks and time of intruder exploration for CRF, UCN3 and OXT-A are presented in Figures 2 and 4; to provide a complete overview of neuropeptidergic contribution to maternal aggression, the same data are reported here.

| Group | Occurrence<br>[n] | Time<br>[s] |  |  |  |  |
| --- | --- | --- | --- | --- | --- | --- |
|  | Attack | Attack | Pinning | Intruder<br>exploration | Self-grooming | Resting |
| VEH | 4.8 ± 1.5 | 3.3 ± 1.1 | 12.8 ± 4.9 | 107.7 ± 18.1 | 68.9 ± 11.8 | 22.0 ± 10.8 |
| CRF | **1.0 ± 0.3 #** | 0.8 ± 0.3 | 4.7 ± 3.4 | 102.8 ± 16.4 | **126.8 ± 22.1 #** | 22.0 ± 6.6 |
| VEH | 12.7 ± 2.0 | 11.2 ± 2.2 | 64.0 ± 8.9 | 90.7 ± 13.8 | 53.5 ± 9.2 | 6.1 ± 2.9 |
| CP-154,526 | 16.9 ± 2.9 | 13.1 ± 2.7 | **130.0 ± 22.8 #** | 62.0 ± 9.8 | 40.6 ± 9.5 | 1.2 ± 0.8 |
| VEH | 7.6 ± 2.3 | 7.5 ± 3.0 | 59.0 ± 20.5 | 45.4 ± 13.5 | 68.8 ± 20.0 | 4.0 ± 2.2 |
| CP-154,526 + CRF | 4.2 ± 1.6 | 3.8 ± 1.2 | 16.8 ± 5.9 | 74.5 ± 14.8 | 80.8 ± 13.6 | 12.2 ± 4.2 |
| VEH | 8.1 ± 2.6 | 8.7 ± 2.8 | 62.4 ± 25.0 | 65.5 ± 21.4 | 85.5 ± 20.4 | 1.0 ± 0.7 |
| UCN3 | **1.8 ± 0.6 #** | **1.1 ± 0.3 #** | **12.3 ± 5.8 (#)** | 98.4 ± 11.9 | 86.3 ± 17.7 | **55.7 ± 18.6 ##** |
| VEH | 4.0 ± 1.4 | 4.7 ± 1.6 | 29.4 ± 8.9 | 97.0 ± 17.4 | 89.0 ± 17.7 | **8.0 ± 3.5 (#)** |
| Astressin 2B | 10.1 ± 2.9 | 8.2 ± 2.7 | 60.1 ± 18.9 | 94.6 ± 15.3 | 52.3 ± 12.6 | 0 ± 0 |
| VEH | 11.6 ± 4.1 | 10.8 ± 4.1 | 64.3 ± 21.9 | 88.1 ± 28.8 | 83.5 ± 28.4 | 1.2 ± 0.8 |
| CRF(6-33) | 9.6 ± 4.2 | 8.8 ± 4.5 | 93.1 ± 43.5 | 118.9 ± 56.7 | **16.6 ± 2.5 (#)** | 3.8 ± 2.8 |
| VEH | 12.1 ± 3.0 | 11.8 ± 3.5 | 101.1 ± 29.7 | 64.4 ± 16.6 | 80.9 ± 24.2 | 0 ± 0 |
| OXT-A | **5.0 ± 1.3 (#)** | **4.1 ± 1.3 (#)** | 77.2 ± 27.0 | **170.7 ± 38.8 #** | 80.4 ± 13.1 | 0 ± 0 |

Next, we correlated the number of attacks with the occurrence of nursing measured at time 60 after the MDT (t60; Figure 2G). We found a significant positive correlation, i.e., less aggressive dams cared less for the pups following stress exposure and *vice versa* (r = 0.603, p < 0.05; Spearman’s correlation). Indeed, acute infusion of CRF also impaired maternal behaviour after the MDT; analysis of total nursing (Figure 2H) revealed a significant main effect of time (F[2, 38] = 9.44, p < 0.01, η^2^ = 0.33; 2-way repeated measures ANOVA), treatment (F[1, 24] = 22.4, p < 0.0001, η^2^ = 0.48) and their interaction (F[2, 48] = 7.87, p < 0.01, η^2^ = 0.25). In detail, dams receiving CRF showed a significant reduction in nursing compared to their respective basal level at both observation times (t30: p < 0.001; t60: p < 0.05 vs basal; Bonferroni post hoc comparisons). Furthermore, CRF-treated rats displayed less nursing compared to VEH-treated rats at both observation times (p < 0.001). This reduction was paralleled by a significant increase of self-grooming after the MDT induced by CRF infusion (time: F[2, 43] = 9.97, p < 0.001, η^2^ = 0.32; treatment: F[1, 24] = 22.9, p < 0.0001, η^2^ = 0.49; interaction: F[2, 48] = 5.11, p < 0.01, η^2^ = 0.18; Figure 2I). At t30 after infusion, CRF-treated dams showed significantly more self-grooming both compared to basal and to VEH-treated rats (p < 0.001). This effect was almost back to basal levels at t60 after infusions, as statistical analysis revealed only a trend towards higher self-grooming compared to basal values (p = 0.059).

The acute inhibition of the CRF-BP via CRF(6-33) infusion, which should result in increased endogenous CRF levels and, thus, heightened CRF-R signalling (Ketchesin et al., 2017), did not affect maternal aggression or maternal care (Table 3 and Supplementary Figure S1A; see Supplementary Tables S1 and S2 for statistical details of maternal care).

#### 3.1.2. CRF-R1 inhibition increased defensive aggressive behaviour

The acute infusion of the selective CRF-R1 antagonist CP-154,526 induced a significant increase of time spent on pinning down the intruder (t[13] = 3, p < 0.05, Cohen’s *d* = 1.16; Table 3), a defensive aggressive behaviour (Bosch, 2013). All other aggressive and non-aggressive behaviours were not affected by this treatment. We anticipated that exogenous CRF infusion would activate CRF-R2 in case CRF-R1 are unavailable, thus impairing maternal aggression. Importantly, the double injection of CP-154,526 followed by CRF did not affect maternal aggression (Table 3), suggesting that CRF is mainly acting on CRF-R1 in controlling this aspect of maternal behaviour.

As for maternal care, CP-154,526 did not affect this behaviour under stress conditions (Supplementary Figure S1B; Supplementary Tables S1 and S2 for statistical details). When analysing the effects of the double infusion of CP-154,526 followed by CRF, we found significant differences in total nursing at t30 after stress compared to the basal level (p < 0.001, Bonferroni post-hoc comparisons, Supplementary Figure S1C; Supplementary Table S1 for statistical details), as well as at t60 compared to t30 (p<0.01), regardless of the treatment. Furthermore, analysis of self-grooming (Supplementary Figure S1C; Supplementary Table S2 for statistical details) revealed a significant difference in both groups at t30 compared to basal (p < 0.01, Bonferroni multiple comparisons), as well as at t60 compared to t30 (p<0.001).

#### 3.1.3. CRF-R2 activation impaired maternal aggression without affecting care

The experimental timeline is shown in Figure 3A. The representative ethograms of one VEH- and one UCN3-treated mother indicate that UCN3 infusion reduced the occurrence of aggressive behaviours during the MDT (Figure 3B). Indeed, CRF-R2 activation via UCN3 infusion significantly decreased the number of attacks (U = 8, p ≤ 0.05, Mann Whitney test; Cohen’s *d* = 1.2; Figure 3C) and increased the latency to first attack (U = 2, p < 0.01, Mann Whitney test; Cohen’s *d* = 1.3 Figure 3D). Moreover, the overall z-score for aggression was significantly reduced compared to VEH-treated rats (t(13) = 4.49, p < 0.001, unpaired t-test; Figure 3E). Regarding non-aggressive behaviours, the time spent exploring the intruder was not altered by the treatment (p > 0.05, Figure 3F), while the time spent resting was increased (p < 0.01 vs VEH-treated rats, Cohen’s *d* = 1.6; Table 3). However, maternal care after the MDT was not altered by selective CRF-R2 activation (Supplementary Figure S2A; Supplementary Tables S1 and S2 for statistical details). Lastly, the inhibition of CRF-R2 via Astressin 2B infusion did not alter maternal aggression or care after stress (Table 3 and Supplementary Figure S2B; see also Supplementary Tables S1 and S2).

**Figure 3.**
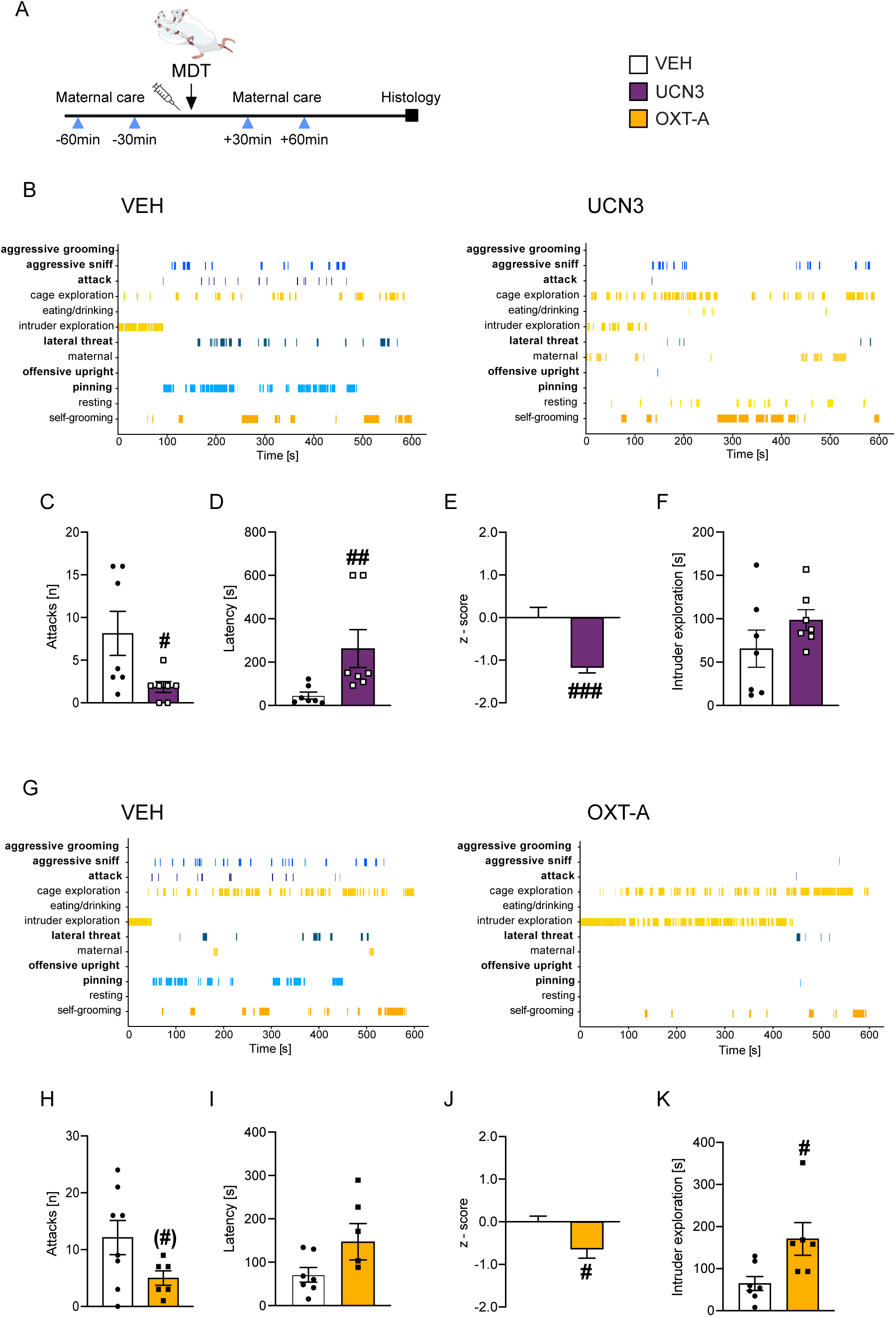
Infusion of UCN3 or of OXT-A in the NAcSh impaired maternal aggression in lactating rats. (A) Experimental timeline (for details, see main text, paragraph 2.2.1.). (B, G) Representative behavioural ethograms during the MDT of one dam, each, treated with (B) vehicle (VEH) or UCN3, or (G) VEH or OXT-A; aggressive behaviours are highlighted in bold and represented by blue bars. Behavioural parameters measured during the MDT: (C, H) number of attacks, (D, I) latency to first attack, (E, J) overall z-score for aggression, (F, K) time spent exploring the intruder. (#) p ≤ 0.07, # p ≤ 0.05, ## p < 0.01, ### p < 0.001 vs VEH-treated dams, Unpaired t test, or Welch’s t test or Mann-Whitney test.

#### 3.1.4. OXT-R inhibition impaired maternal aggression without affecting care

The representative ethograms (Figure 3G) of one VEH- and one OXT-A-treated mother highlight that OXT-A infusion impaired maternal aggression during the MDT. OXT-A induced a trend towards reduced attacks (t[9] = 2.18, p = 0.056, Welch’s t-test; Cohen’s *d* = 1.1; Figure 3H), without affecting the latency to first attack (p > 0.05; Figure 3I). However, we found a significant reduction of the overall z-score for aggression (t[11] = 2.64, p < 0.05, unpaired t-test; Figure 3J). Lastly, dams treated with the OXT-R antagonist showed increased time spent exploring the intruder (U = 4, p < 0.05, Mann Whitney test; Cohen’s *d* = 1.4; Figure 3K). However, the treatment with OXT-A did not induce any effect on maternal care under stress conditions (Supplementary Figure S2C; Supplementary Tables S1 and S2 for statistical details).

### 3.2. Descriptive approach

#### 3.2.1. MDT exposure increased activation of CRF+ cells within the NAcSh

Considering the strong effects on maternal aggression induced by CRF-R and OXT-R pharmacological manipulation, we investigated the mRNA expression of *Crh*, *Crh-r1*, *Crh-r2*, *Crh-bp* and *Oxt-r* in the NAcSh of lactating rats with / without exposure to the MDT combined with / without the presence of the pups (Figure 4A). No changes in mRNA levels were observed for any of the tested genes in either of the groups (Figure 4B-F; for statistical analysis, see Supplementary Table S3). However, in the *MDT - No pups* group, *Crh-bp* mRNA levels positively correlated with the number of pinning down (r = 0.81; p = 0.027; R^2^ = 0.66; Pearson’s correlation; Supplementary Figure S3), while no further correlations with maternal aggression or care were found (data not shown).

**Figure 4.**
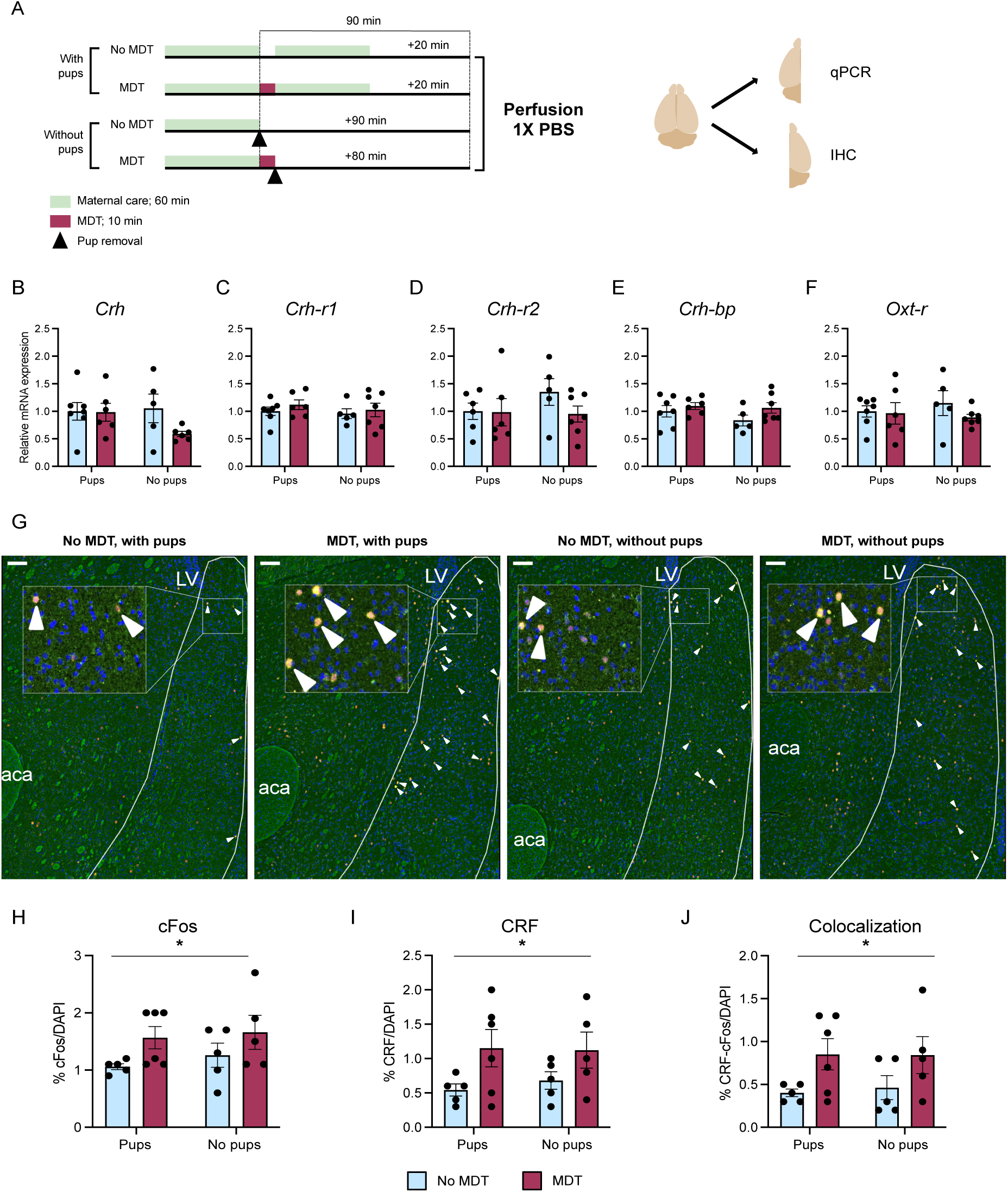
MDT exposure increased CRF+ cells activation in the NAcSh of lactating rats. (A) Overview of the experimental designs (for details, see main text 2.2.2., Experiment 2); brain icon adapted from (Shin), SciDraw.io. mRNA levels of (B) *Crh*, (C) *Crh-r1*, (D) *Crh-r2*, (E) *Crh-bp* and (F) *Oxt-r* in punches of the whole NAc of lactating rats with / without exposure to the MDT and with / without the presence of the pups. Data are expressed as relative levels vs control group (*No MDT - with pups*) and are presented as mean ± SEM. (G) Representative fluorescent images showing colocalization (yellow) of c-Fos (red) / CRF (green) / DAPI (blue) immunoreactive cells in the NAcSh comparing all four groups (scale bar 100 µm; white arrows indicating representative colocalizing cells). Percentage of cells labelled with (H) cFos, (I) CRF, and (J) their co-labelling. Data are presented as mean ± SEM. * p ≤ 0.05, main effect of MDT, 2-way ANOVA. Abbreviations: LV, lateral ventricle; aca, anterior part of the anterior commissure.

Furthermore, we performed fluorescent IHC to visualise cFos and CRF positive cells (for representative colocalization staining, see Figure 4G). 2-way ANOVA analysis of cFos+ cells revealed a significant main effect of the MDT (F[1, 17] = 4.7, p < 0.05; Figure 4H), but not of the pups’ presence (F[1, 17] = 0.493, p > 0.05) or their interaction (F[1, 17] = 0.065, p > 0.05). Similarly, we found a significant main effect of the MDT on the percentage of CRF+ cells (F[1, 17] = 6, p = 0.026; Figure 4I) but neither of the presence of the pups (F[1, 17] = 0.066, p > 0.05), nor of their interaction (F[1, 17] = 0.157, p > 0.05). Lastly, the statistical analysis of the % of cells expressing both cFos and CRF revealed a significant main effect of the MDT (F[1, 17] = 6.5, p = 0.021; Figure 4J), while no effects of pups’ presence (F[1, 17] = 0.023, p > 0.05) or the interaction of the factors (F[1, 17] = 0.046, p > 0.05) were found. Lastly, no correlations between levels of cFos, CRF or their colocalization and maternal aggression or care were observed (data not shown).

#### 3.2.2. Exposure to the MDT elicited OXT release in the NAc

To further characterize the neurobiological changes in response to the MDT, we measured OXT release before, during and after the encounter with an unfamiliar intruder (Figure 5A). 1-way repeated measures ANOVA analysis did not reveal any significant effect of exposure to the MDT (F[2, 12] = 2.2, p = 0.15, η^2^ = 0.27; Figure 5B). However, when analysing the OXT content during the MDT compared to the average of the two basal samples (Figure 5C), we found an increase in OXT levels in response to the MDT, despite not reaching statistical significance (t[5] = 2.3, p = 0.07, Paired t test; Cohen’s *d* = 2). Furthermore, we found a negative correlation between the SON *Oxt* mRNA levels and the release of OXT in the NAc (r = −0.853, p < 0.05, R^2^ = 0.73; Pearson’s correlation; Figure 5D), while no correlation was found between PVN *Oxt* mRNA levels and OXT release in the NAc (data not shown). Neither OXT release nor *Oxt* mRNA levels in the PVN or SON correlated with aggressive behaviours during the MDT (data not shown).

**Figure 5.**
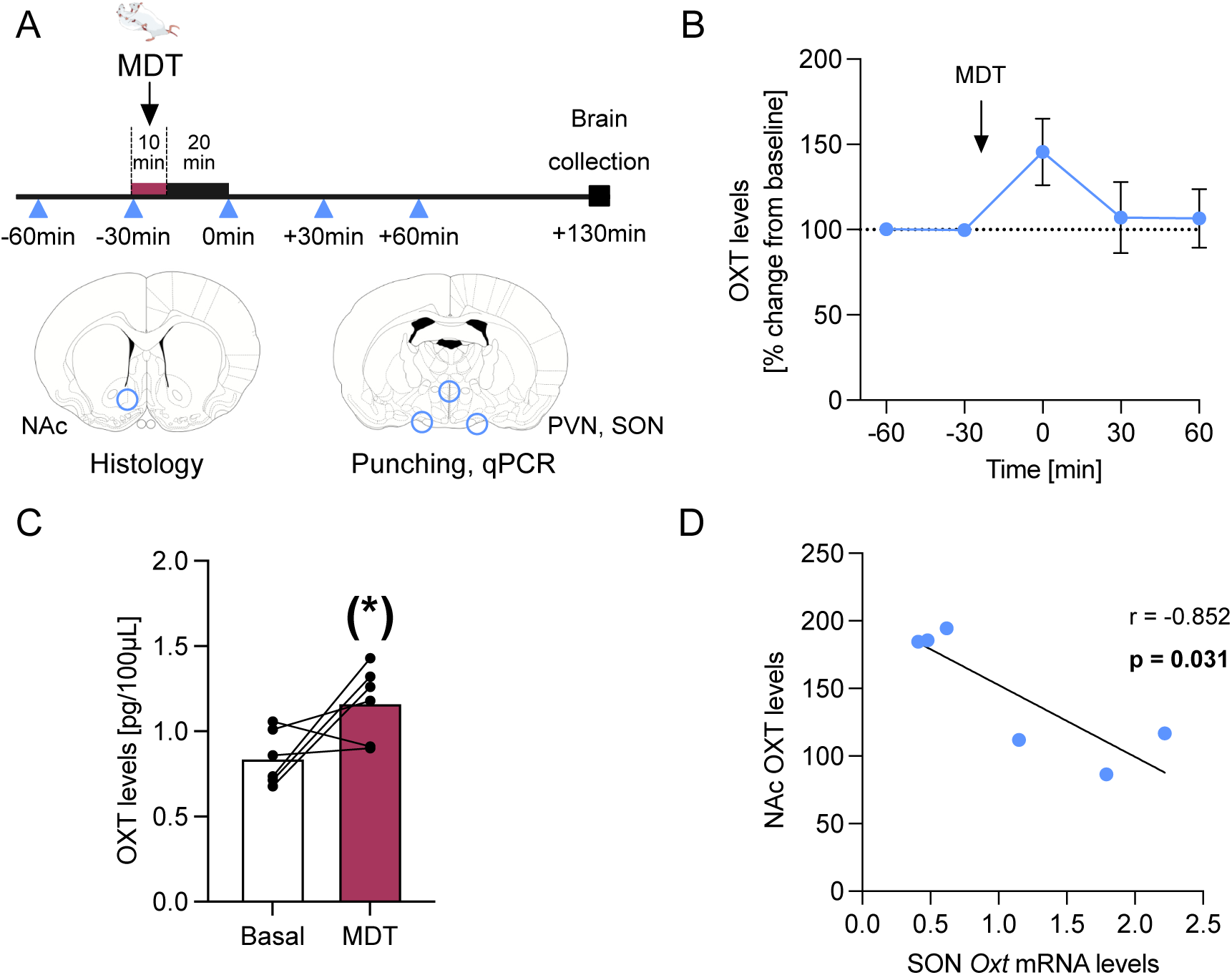
MDT exposure induced OXT release in the NAc. (A) Overview of the experimental timeline (for details see main text, paragraph 2.4.3). The red line marks the duration of the MDT; the blue triangles mark the collection of microdialysate samples. Brain coronal sections adapted from (Paxinos and Watson, 2013). (B) Percentage change of intra-NAc OXT release, and (C) OXT content compared between mean basal samples and in response to the MDT. (D) Pearson’s correlation between intra-NAc OXT release and SON *Oxt* mRNA levels. Data are presented as mean ± SEM. (*) p ≤ 0.07 vs basal, Unpaired t-test. Abbreviations: MDT, maternal defence test; NAc, nucleus accumbens; PVN, paraventricular nucleus; SON, supraoptic nucleus.

#### 3.2.3. Intra-NAc retrodialysis of CRF induced local release of OXT and DA

Since exposure to the MDT is a psychosocial challenge that elicits a stress response (Neumann et al., 2001) and increases the release of OXT in certain brain areas (Bosch, 2013) including the NAc (this study; see 3.2.2.), we measured intra-NAc OXT release in response to CRF-R1 or CRF-R2 activation (infusion of CRF or UCN3, respectively; Figure 6A). The mean baseline OXT release was not different between CRF- or UCN3-treated rats prior to retrodialysis (CRF: 1.172 ± 0.21 pg/30µL; UCN3: 1.140 ± 0.1 pg/30µL; U = 24, p > 0.05, Mann Whitney test; Figure 6A insert). Mixed-effects analysis for repeated measures revealed a significant effect of time (χ^2^ (1) = 1.26; F[2, 25] = 10, p = 0.0004, Figure 6A), treatment (F[1, 13] = 5.86, p = 0.031) and the interaction of the two factors (F[4, 45] = 3.29, p = 0.019). In detail, retrodialysis of CRF, but not UCN3, induced a significant transient increase in OXT release during retrodialysis (t0) compared to both basal samples (p < 0.01, Bonferroni’s multiple comparisons).

**Figure 6.**
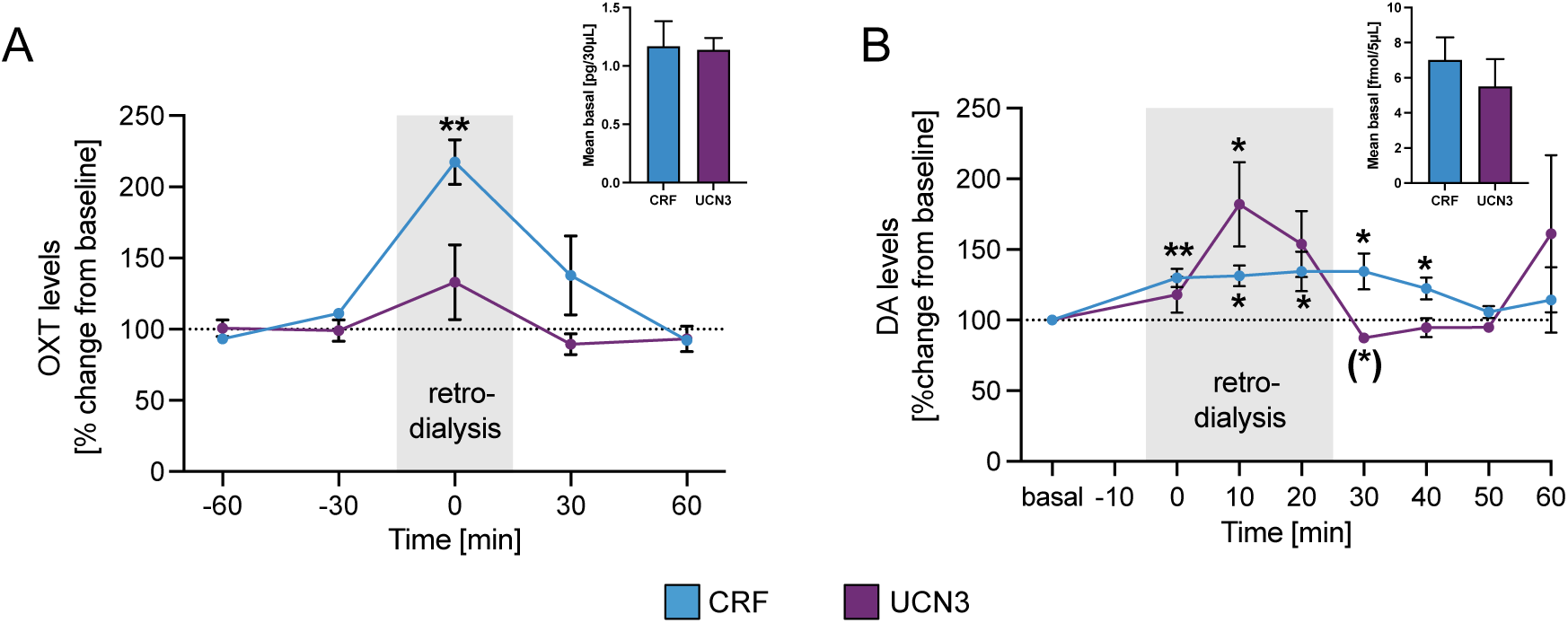
Intra-NAc activation of CRF-R1 induced local OXT and DA release in lactating rats. Effects of intra-NAc retrodialysis of agonists (ago) for CRF-R1 (CRF) or CRF-R2 (UCN3) on local release of (A) OXT (CRF-R1 ago n = 5, CRF-R2 ago n = 10) and (B) DA (CRF-R1 ago n = 8, CRF-R2 ago n = 5). Data are presented as mean ± SEM; (*) p ≤ 0.07, * p ≤ 0.05, ** p < 0.01 vs respective basal; Mixed-effects analysis for repeated measures followed by (A) Bonferroni’s post hoc comparisons or (B) Fisher’s LSD.

We further characterized the interactions between the CRF and the DA systems within the NAc. Therefore, we measured intra-NAc DA release in response to either CRF-R subtype activation (Figure 6B). The mean baseline DA release was not different between CRF- or UCN3-treated rats prior to retrodialysis (CRF: 7.01 ± 1.3 fmol/5µL; UCN3: 5.49 ± 1.6 fmol/5µL; t[11] = 0.74, p > 0.05, Unpaired t test; Figure 6B insert). Statistical analysis over the whole time course revealed a significant effect of time and time x treatment interaction (χ^2^ (1) = 5.3; time: F[3, 28] = 4.9, p < 0.001; time x treatment: F[7, 64] = 2.6, p < 0.05, Mixed-effects analysis for repeated measures; Figure 6B), while no effects were detected for treatment alone (F[1, 11] = 0.30, p > 0.05). In detail, CRF induced a significant and long-lasting release of DA from the start of retrodialysis at t0 (p < 0.01 vs respective basal, Fisher’s LSD) until t40 (p < 0.05 from t10 to t40 vs respective basal). In contrast, retrodialysis of UCN3 induced a sharp, but transient significant increase of DA release at t10 compared to basal (p < 0.05), and tended to be below baseline after termination of UCN3 retrodialysis at t30 (p = 0.057).

#### 3.2.4. *In-situ* hybridization revealed differential distribution and expression patterns of CRF-R subtypes in the NAc

We investigated the neurochemical properties of cells expressing *Crh-r* and their distribution along the rostro-caudal axis of the NAc in lactating rats using ISH. For this, either *Crh-r1* or *Crh-r2* probes were used in combination with *Ppp1r1b* and *Pvalb* probes, labelling MSN and GABAergic interneurons, respectively (for a representative overview, see Figure 7A and B). As depicted in Figure 7C, most *Crh-r1* expressing neurons were located in the central part of the NAc (NAcCo: 42,23 %; NAcSh: 41,86 %), whereas the least expression was observed in the rostral pole (NAcCo: 26,03 %; NAcSh: 22,85 %), irrespective of NAcCo or NAcSh subdivision (% of all *Crh-r1* colocalized with DAPI, regardless of colocalization with *Ppp1r1b* and / or *Pvalb*). The highest percentage of colocalization of *Crh-r2* with DAPI was found towards the rostral pole of the NAcCo (35,46 %) and NAcSh (38,03 %) (see Figure 7D), while the lowest expression was measured in the caudal pole (NAcCo: 30,33 %, NAcSh: 25,53 %). However, the overall distribution of either *Crh-r1* or *Crh-r2* in the NAcCo compared to the NAcSh was proportionally similar (Figure 7C and D).

**Figure 7.**
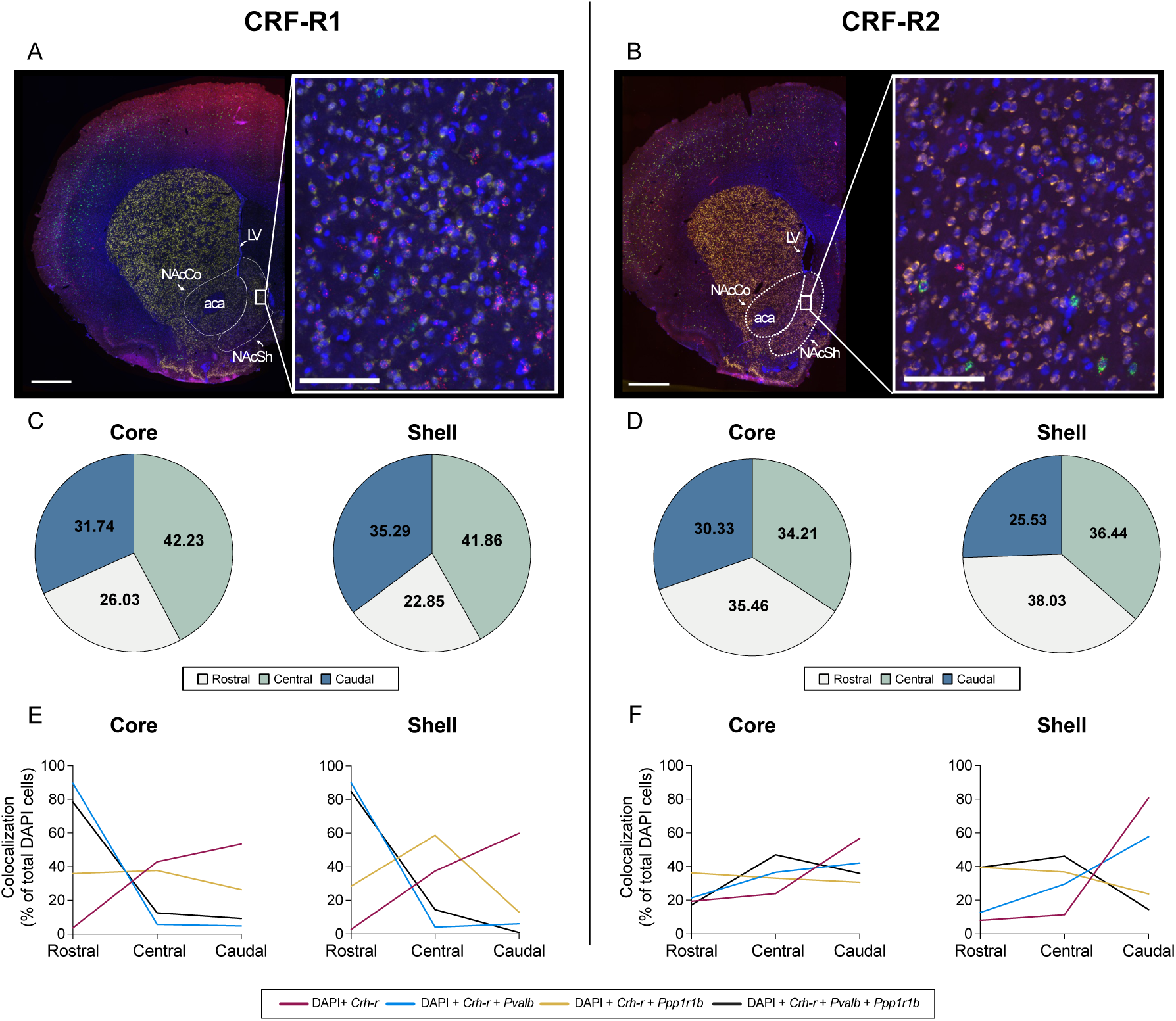
Neurochemical properties and distribution pattern of *Crh-r1* and *Crh-r2* along the rostro-caudal axis of the NAcCo and NAcSh. Results of ISH within the NAc of lactating rats for *Crh-r1* (left) and *Crh-r2* (right). ISH representative picture of the central NAc plane for (A) *Crh-r1* or (B) *Crh-r2* (n = 3; scale bar: 1000 µm). Close-up pictures of the NAcCo and NAcSh reveal colocalization of DAPI (blue) with either *Crh-r* (red), *Ppp1r1b* (yellow), *Pvalb* (green), or all markers (scale bar: 100 µm; white rectangles indicate the close-up area; white dashed lines indicate the NAc region). Distribution of (C) *Crh-r1* in the NAcCo (left) and NAcSh (right), (D) *Crh-r2* in the NAcCo (left) and NAcSh (right). Pie chart data are expressed as % compared to the whole NAc for all rats (C, D; for details see 2.7.3.). Neurochemical properties of (E) *Crh-r1* in the NAcCo (left) and NAcSh (right), (F) *Crh-r2* in the NAcCo (left) and NAcSh (right). Data are presented as % of NAc cells expressing a specific marker in each plane on the total count of each marker for the whole NAc (E, F; for details see 2.7.3.). Abbreviations: aca, anterior part of the anterior commissure; LV, lateral ventricle; NAcCo, nucleus accumbens core; NAcSh, nucleus accumbens shell.

Furthermore, *Crh-r1* in the rostral portion of the NAcCo and NAcSh were mostly colocalized with *Pvalb* (NAcCo: 89,5 %; NAcSh: 90 %) (see Figure 7E). The % of *Crh-r1*+ neurons colocalizing with *Ppp1r1b* was the highest in the central plane (*Ppp1r1b,* NAcCo: 37,7 %; NAcSh: 58,7 %) and decreased towards the rostral and caudal plane of both NAcCo and NAcSh. The highest expression of *Crh-r1* in combination with both *Ppp1r1b* and *Pvalb* was observed in the rostral pole (NacCo: 78.3 %; NAcSh: 84.7 %) and decreased towards the caudal pole, while the % of DAPI cells expressing only *Crh-r1* was the highest in the caudal part of the NAcCo (42.9 %) and NAcSh (37.4 %). On the other hand, colocalization of *Crh-r2* and *Pvalb* increased from the rostral (NAcCo: 21,3 %; NAcSh: 12,7 %) to the caudal pole (NAcCo: 42,1 %, NAcSh: 57,8 %) both in the NAcCo and in the NAcSh (Figure 7F), while colocalization of *Crh-r2* with *Ppp1r1b* decreased in this direction, specifically in the NAcSh. In contrast to *Crh-r1*, expression of *Crh-r2* in combination with both *Ppp1r1b* and *Pvalb* was the highest in the central plane (NAcCo: 46.9 %; NAcSh: 46.1 %). Lastly, the % of *Crh-r2* cells expressing only this receptor was the highest in the caudal part of the NAcCo (56.8 %) and NAcSh (80.7 %).

#### 3.2.5. Retrograde tracing revealed sources of CRF+ neurons in NAcSh

To identify major sources of CRF+ neurons in the NAcSh, virgin female rats were infused with a retrogradely transported tracer into the NAcSh (Figure 8A). We identified the basolateral amygdala, part of the medial prefrontal cortex (mPFC, including prelimbic and infralimbic cortex), the paraventricular thalamus (PV), dorsolateral entorhinal cortex, perirhinal cortex and ventral subiculum as sources of CRF+ neurons projecting to the NAcSh (Figure 8B and C).

**Figure 8.**
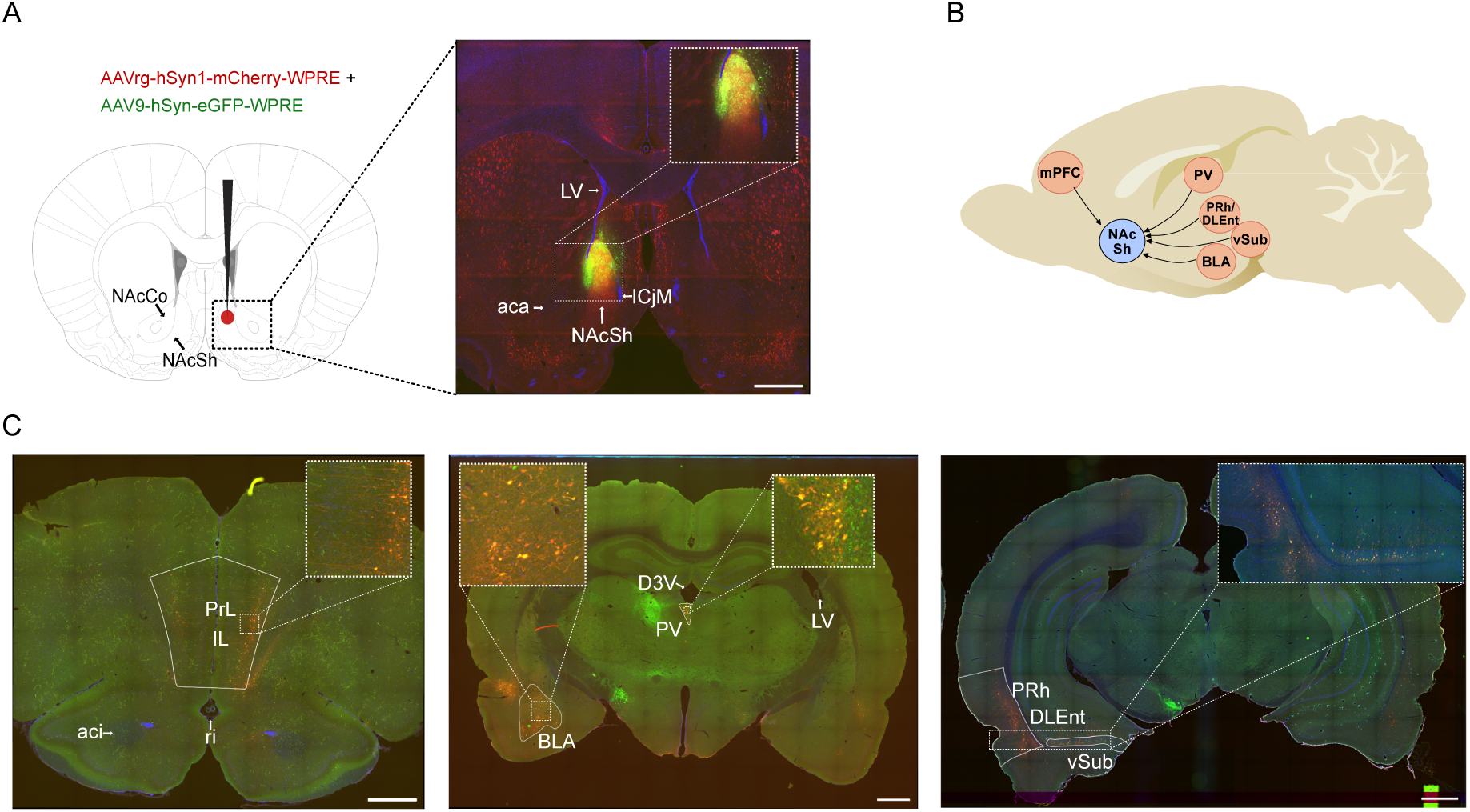
Brain regions with CRF+ neurons innervating the NAcSh in female rats. (A) Representative picture showing the injection site and the corresponding coronal section as well as the injected viruses (from (Paxinos and Watson, 2013)). (B) Schematic overview of CRF-enriched regions innervating the NAcSh; brain icon adapted from (Dooley), SciDraw.io. (C) Representative coronal pictures depicting CRF-enriched regions projecting to the NAcSh (n=2; scale bar: 1000 µm). Abbreviations: aca, anterior part of the anterior commissure; aci, intrabulbar part of the anterior commissure; BLA, basolateral amygdala; DLEnt, dorsolateral entorhinal cortex; D3V, dorsal 3^rd^ ventricle; ICjM, major island of the islands of Calleja; IL, infralimbic cortex; LV, lateral ventricle; NAcCo, nucleus accumbens core; NAcSh, nucleus accumbens shell; MO, medial orbital cortex; mPFC, medial prefrontal cortex; PrL, prelimbic cortex; PV, paraventricular thalamus; PRh, perirhinal cortex; ri, rhinal incisure; vSub, ventral subiculum

## 4. Discussion

In this study, we not only identified the NAcSh as a crucial regulatory brain region for maternal aggression but also demonstrated the specific roles of local neuropeptidergic transmission in the control of this behavioural domain. In detail, intra-NAcSh activation of CRF-R1 via CRF infusion had a modulatory function in controlling maternal aggression as well as aspects of maternal care under those stressful conditions. Local CRF-R2 and OXT-R signalling predominantly modulated active maternal responses, such as pup defence. Although mRNA levels of members of CRF and OXT systems were unchanged, MDT exposure significantly increased the number of cFos-activated CRF+ neurons. Furthermore, local OXT release was increased in dams exposed to the MDT, presumably due to the increased stress level. In that line, we discovered local interactions between the CRF and the OXT systems, as well as the DA system: intra-NAc OXT release increased with infusion of CRF, but not UCN3. Furthermore, local DA levels were enhanced by activation of both CRF-R but in distinct release patters. Regarding the neurochemical properties of *Crh-r*-expressing cells in the NAc and their spatial distribution, *Crh-r1* expression was more abundant in the medial NAc, regardless of NAcCo or NAcSh subdivision, while *Crh-r2* was mostly expressed in the rostral NAc. The co-expression of *Crh-r* and the other markers tested for differed along the rostro-caudal plane. Lastly, we further characterized the mPFC, BLA and PV as main sources of CRF to the NAcSh.

### 4.1. Behavioural effects of neuropharmacological manipulations of the NAcSh CRF and OXT systems

The NAc is part of the reward circuitry (Floresco, 2015; Lewis et al., 2021; Salgado and Kaplitt, 2015), and previous studies defined aggression as potentially rewarding in male mice (Couppis and Kennedy, 2008; Golden et al., 2019), abnormally aggressive male rats (Beiderbeck et al., 2012) but also in female rats (Borchers et al., 2023). Indeed, the NAc is also part of the neurocircuit of aggression, being a region that triggers this behaviour in mice (Aleyasin et al., 2018). In this light, our findings suggest that within the NAcSh both the CRF and OXT systems reduce the reinforcing properties of aggression, since mothers treated with CRF-R agonists (CRF and UCN3) or OXT-A showed dampened aggressive responses towards the intruder (Figures 2 and 3). Strengthening this hypothesis, neuropeptidergic manipulations in the NAcSh specifically reduced aggression and pup-defence, without affecting social behaviour in general. Indeed, the time exploring the intruder was not lower compared to VEH-treated rats, emphasizing that the dams were aware of and interested in the presence of the intruder, thus further excluding a deficit in social interactions as well as a sedation effect. Interestingly, the NAc innervates the lateral habenula (LHb) with inhibitory GABAergic projections; upon stimulation of this inhibitory pathway, the LHb promotes aggression-seeking behaviour, whereas LHb neurons are strongly activated in non-aggressive mice (Aleyasin et al., 2018; Flanigan et al., 2017; Golden et al., 2016). Therefore, CRF-R activation or OXT-R inhibition in the NAcSh might downregulate this GABAergic inhibitory control on the LHb (Aleyasin et al., 2018; Flanigan et al., 2017; Golden et al., 2016), increasing the activity of the LHb neurons and reducing aggressive responses towards the intruder. However, further studies are needed to confirm this modulatory pathway following neuropeptidergic manipulations in lactating rats.

Another interpretation of our findings is that finely tuned neuropeptide transmission in the NAcSh, which integrates stress inputs (Itoga et al., 2019), is in fact essential to trigger maternal aggression. When the baseline stress response is enhanced, i.e., due to increased CRF-R or reduced OXT-R signalling, this prevents the mother from actively defending the offspring (Bosch, 2013). In support, lactating mice show reduced maternal aggression after acute exposure to restraint stress (Gammie and Stevenson, 2006). Besides reducing the number of attacks, CRF infusion increased self-grooming behaviour during the MDT (Figure 2), indicative of stress-induced arousal (Kalueff et al., 2016). Thus, the maternal response to the stressor was shifted from active (defence of the pups) to a more passive and stereotypic response, i.e., self-grooming (Kalueff et al., 2016). Interestingly, increased self-grooming continued after termination of the stress in the present study, and we observed a similar response after CRF infusion under non-stress conditions (Sanson et al., 2024b), which is further in line with a study showing long-lasting behavioural arousal following intra-NAcSh CRF infusion in male rats (Holahan et al., 1997). Previously, we found a similar increase of self-grooming after CRF infusion in the MPOA (Klampfl et al., 2018), suggesting that, overall, CRF shifts the mother’s behaviour from pup care to self-care. In the same line, local CRF infusion in the NAcSh induced a major reduction of nursing behaviour after stress exposure as we saw for the MPOA (Klampfl et al., 2018).

Interestingly, dams receiving intra-NAcSh CRF showed a positive correlation between the number of attacks and the occurrence of total nursing 60 min after the termination of the stress, indicating that more passive mothers (i.e., did not attack) also showed less pup care after stress exposure. This implicates that an acute overstimulation of the CRF-R1 is sufficient to impair different aspects of maternal behaviour long-lastingly. To further strengthen a principal role of CRF-R1 in maternal aggression (Gammie et al., 2004; Klampfl and Bosch, 2019a), acute CRF-R1 inhibition increased defensive aggressive behaviour (pinning down the intruder), while the additional single infusion of CRF had no further effects on maternal aggression or care (Table 3 and Supplementary Figure S1). This proves that CRF mainly acts on CRF-R1 in the NAcSh to acutely regulate different components of maternal behaviour. Furthermore, it indicates that a finely balanced CRF-R1 transmission is necessary for the required display of aggressive behaviours.

Selective CRF-R2 activation via UCN3 as well as OXT-R inhibition strongly impaired maternal aggression (Figure 3), without subsequently affecting maternal care (Supplementary Figure S2). This suggests that a functional transmission of these two receptors in the NAcSh is particularly important during active maternal behaviours, such as pup-defence. Indeed, intact OXT-R transmission has been known to mediate maternal aggression in several brain regions (for review, see (Bosch, 2013)). Furthermore, several studies link hyperactive central CRF-R2 transmission to reduced maternal aggression (D’Anna and Gammie, 2009; D’Anna et al., 2005; Gammie et al., 2008; Gammie et al., 2004) (for review, see (Sanson et al., 2024a)), while sparse evidence has proven its involvement in maternal care so far, for instance in the BNST (Klampfl et al., 2014, 2016). However, no effects of acute CRF-R2 inhibition via Astressin 2B infusion in the NAcSh were found on any aspects of maternal behaviour (Table 3 and Supplementary Figure S2), suggesting that endogenous CRF mostly activates the CRF-R1 during maternal aggression.

### 4.2. Effects of MDT exposure on CRF and OXT systems mRNA in the NAc

Given the stressful nature of the MDT (Bosch, 2013; Gammie et al., 2008; Neumann et al., 2001), we expected acute changes in gene expression of members of the CRF and OXT systems; however, their mRNA expression remained unchanged (Figure 4). Interestingly, *Crh-bp* mRNA levels positively correlated with the number of pinning during the MDT only in the *MDT - No pups* group, perhaps indicating a compensatory mechanism to reduce the stress response to pup removal in mothers that were more protective (Supplementary Figure S3). However, as the mRNA was isolated from punches of the whole NAc, we cannot exclude that we failed to detect further subtle changes within the subregions of the NAc itself in response to the psychosocial stressor. Indeed, the analysis of protein levels in the NAcSh provided a significant main effect of the MDT on cells expressing cFos, CRF, and their double labelling (Figure 4 G-J). Following the stressful MDT, the activity of CRF+ cells in the NAcSh of lactating rat mothers increased regardless of pup presence. This is supported by previous findings in the BNST and PVN where acute stress exposure increased Fos-CRF colocalization (Fetterly et al., 2019; Walker et al., 2019). Strengthening our results, the NAcSh is more activated than the NAcCo during aggression seeking in male mice, as indicated by higher Fos expression (Golden et al., 2019). Taken together, a certain level of CRF in the NAcSh is necessary to adequately respond to a social threat. However, since the NAcSh receives CRF inputs from several stress-related brain regions (Itoga et al., 2019), it is crucial to measure CRF release during the MDT within the NAcSh to fully characterize the involvement of the local CRF systems in maternal aggression.

### 4.3. Effects of MDT exposure on OXT release in the NAc

Interestingly, the levels of OXT in the NAc tended to increase during exposure to the MDT (Figure 5), similar to what we described for the PVN, central amygdala and BNST (Bosch et al., 2004; Bosch et al., 2005), suggesting that this neuropeptide is a crucial mediator of pup defence in different brain regions. The intra-NAc OXT levels were negatively correlated with SON *Oxt* mRNA expression. We speculate that the activated OXT system in the NAc during exposure to the MDT subsequently inhibits OXT producing cells in the SON, in a sort of negative feedback loop. This hypothesis is based on a previous study demonstrating that OXTergic cells in the SON get inhibited after NAc stimulation (Shibuki, 1984). Since the SON is thought to be mostly involved in the peripheral release of OXT (Jurek and Neumann, 2018; Menon and Neumann, 2023), it might be dampened by exposure to the MDT, as most OXT release might be redirected to the brain to promote the active defence of the offspring. Furthermore, this might be a physiological response to the lack of milk ejection reflex during MDT, when the mother does not nurse the offspring. However, additional studies are needed to characterize such a potential modulation.

### 4.4. Effects of intra-NAc CRF-R activation on local OXT and DA release

As known for other brain regions, the CRF system can interact with the OXT system (Bosch et al., 2016; Dabrowska et al., 2011; Klampfl et al., 2018; Martinon and Dabrowska, 2018; Winter and Jurek, 2019) and the DA system (Chen et al., 2012; Kelly and Fudge, 2018; Lemos et al., 2012), highlighting finely balanced mechanisms to control complex behavioural responses. Expanding this knowledge, we found a CRF-R subtype-specific interaction with the OXT system within the NAc of lactating rats: local infusion of CRF transiently increased OXT release (Figure 6A). A similar mechanism has been shown for the MPOA of lactating rats (Klampfl et al., 2018) as well as the BNST of male rats (Martinon and Dabrowska, 2018). Moreover, we could demonstrate that the activation of either CRF-R induced a significant release of DA within the NAc of lactating rats (Figure 6B) similar to male mice (Lemos et al., 2012) and male rats (Chen et al., 2012). Interestingly, the DA release pattern depended on the activated CRF-R subtype; increased CRF-R1 signalling via CRF caused a sustained release of DA for the duration of CRF retrodialysis and up to 20 min afterwards, whereas CRF-R2 activation induced a sharp, transient increase of DA release. We hypothesize that the different nature of CRF-R1- vs CRF-R2-expressing cells in the medial NAc (Figure 7) might contribute to the distinct release patterns. Remarkably, DA release pattern in the NAc is long-lasting in response to reward-related cues, whereas a transient peak occurs for avoidance-related cues, such as a shock (Gentry et al., 2016). Hence, our findings confirm in lactating rats what was found in male mice, i.e., that acute CRF infusion represents an appetitive stimulus (Lemos et al., 2012), and suggest that CRF-R2 activation via UCN3 retrodialysis might represent a rather aversive stimulus for the maternal brain. The increased release of OXT and DA might also represent a compensatory mechanism put in place by the lactating brain to counteract the detrimental effects of hyperactive CRF-R1 transmission on maternal behaviour. However, such mechanisms might not be sufficient to fully rescue the negative effects of CRF-R1 hyperactivation when CRF levels exceed the physiological range, e.g., due to exposure to severe stress, given that CRF can also elicit acetylcholine release, inducing aversive responses (Chen et al., 2012). Furthermore, the release patterns of OXT and DA upon CRF-R stimulation might be different in response to the MDT, thus further studies should investigate this aspect. Considering the rewarding nature of aggression (Beiderbeck et al., 2012; Borchers et al., 2023; Couppis and Kennedy, 2008; Golden et al., 2019), a certain amount of CRF-R signalling is necessary to elicit DA release (Chen et al., 2012; Lemos et al., 2012). This release might be necessary to process the cues coming from the intruder as appetitive, while excessive activation of the system might impair this pathway. The same might be true for OXT release, as OXT can also modulate reward-related behaviours (Baskerville and Douglas, 2010; Love, 2014). However, the neuroendocrine modulation of maternal aggression differs from other kinds of aggression (for a review, see (Trainor et al., 2009)), thus the rewarding properties of maternal aggression should be further investigated.

### 4.5. Neurochemical properties and distribution of Crh-r expressing cells in the NAc

Several studies postulate a neurochemical or receptor gradient that differs along the rostro-caudal axis of the rat NAc (Mathieu et al., 1996; Ranaldi and Beninger, 1994; Rogard et al., 1993; Voorn and Docter, 1992). Here, we characterized the neurochemical properties and distribution of *Crh-r* expressing cells throughout the NAc of lactating rats. *Crh-r1* were mostly expressed in the central part of the NAcCo and NAcSh, while their expression was reduced towards the rostral and caudal poles of both NAc subregions, with the lowest expression measured at the rostral pole (Figure 7C). *Crh-r2* expression decreased from the rostral to the caudal pole of the NAcCo and NAcSh in lactating rats (Figure 7D). Interestingly, the distribution of *Crh-r1* and of *Crh-r2* was different along the rostro-caudal axis of the NAc. A previous study showed opposite motivational responses following GABA_A_ activation in the rostral or caudal NAcSh, implying that the NAcSh is functionally heterogeneous (Reynolds and Berridge, 2001). Thus, our data suggest distinct functions of the CRF-R subtypes in this brain region based on their location, although further studies are warranted to characterize the behavioural responses following CRF-R activation along the rostro-caudal axis. However, both the NAcCo and NAcSh showed a relatively similar distribution of the two receptor subtypes.

In the rostral pole of both the NAcCo and NAcSh, *Crh-r1* were mainly expressed by GABAergic interneurons (Figure 7E). However, this changed in the central and caudal plane of the NAc. Specifically, in both planes of the NAcCo, *Crh-r1* were primarily expressed on cells not expressing any other used markers (*Pvalb* and *Ppp1r1b*). In the central NAcSh, *Crh-r1* were mostly expressed by MSN, the major cell type in the NAc (Castro and Bruchas, 2019; Kardos et al., 2019; Robison and Nestler, 2011), while in the caudal part they were not colocalizing with the other markers. Interestingly, *Crh-r2*-expressing MSN decreased from rostral to caudal (Figure 7F), following the general distribution of *Crh-r2* (as described above), while the portion of *Crh-r 2*expressed on GABAergic interneurons increased. We further found *Crh-r*-expressing cells expressing both *Pvalb* and *Ppp1r1b* in all subregions of the NAc. This finding is in line with a previous study showing that GABAergic interneurons can make synaptic contact onto the soma of MSN in the striatum (Kubota and Kawaguchi, 2000). Overall, our findings confirm sequencing studies in mice revealing a transcriptional gradient of MSN markers in the striatum, including the NAc (Gokce et al., 2016; Saunders et al., 2018). Furthermore, a recent study showed that *Crh-r1* are present on cholinergic interneurons that co-express estrogen receptor alpha in the NAc of male and female mice (Olson et al., 2024). In this light, the cells expressing only the *Crh-r* found in this work might be cholinergic interneurons, although this should be investigated in lactating rats. Adding to the existing literature, the present findings contribute to the understanding of the heterogeneity of the brain CRF system.

### 4.6. Identification of CRF+ projections to the NAcSh

Previous studies identified different innervations to the NAcCo and NAcSh (Brog et al., 1993; Floresco, 2015; Salgado and Kaplitt, 2015). A recent study in male and female mice identified CRF-rich innervations to the NAc from brain regions involved in stress regulation, highlighting the mPFC, BLA and PV as major sources (Itoga et al., 2019). While species differences might exist at the level of the CRF system (Douglas et al., 2003), we confirmed a rich innervation from those regions (Figure 8). In line with Itoga *et al*. (Itoga et al., 2019), we identified part of the mPFC (prelimbic, infralimbic, medial orbital cortex), BLA, PV, ventral subiculum, perirhinal and entorhinal cortex as CRF-enriched projections to the NAcSh. Differently from mice (Itoga et al., 2019), we could not find CRF projections from the BNST, the CA1 region of the hippocampus and the anterior cingulate cortex to the NAcSh in female rats. However, Itoga *et al*. mostly targeted the NAcCo (Itoga et al., 2019), while we focused on the NAcSh. Thus, we cannot exclude that the same regions send CRF innervation to the NAcCo of female rats. In conclusion, sources of CRF+ neurons innervating the NAcSh are mostly conserved across different species, emphasizing the NAc as a crucial region for integrating stress signals.

## 5. Conclusions

The present study provides novel insights into the neurobiological basis of maternal aggression.

To our knowledge, this is the first time linking the NAcSh to maternal aggression, specifically via CRF and OXT systems transmission. Altogether, it highlights novel and complex modulations that contribute to the onset of maternal aggression and defence of the offspring, thereby advancing our understanding of the neuroendocrine basis of such an essential protective behaviour in the postpartum period.

## Acknowledgements

This study was supported by the Deutsche Forschungsgemeinschaft (DFG; BO1958/8-2 to O. J. B.).

We thank M. Fuchs, A. Havasi and R. Maloumby for excellent technical support. We further thank C. Baron, A. L. Boos, N. Hempel, J. Matas Teruel, S. Peña Peña, C. Pérez Gozalbo, L. Reimer, E. Rocaboy and S. Schraml for their helpful assistance. We also thank Dr. D. Slattery, Dr. B. Di Benedetto and Dr. F. Martínez García for the insightful scientific discussions. We express our gratitude to Dr. M. Manning for the supply with oxytocin receptor antagonist.

## CRediT authorship contribution statement

**Alice Sanson:** writing – original draft, review & editing; methodology; investigation; data visualization; formal analysis.

**Annika Köck:** investigation; formal analysis; writing – review & editing

**Luisa Demarchi:** investigation; methodology; writing – review & editing.

**Karl Ebner:** investigation; writing – review & editing.

**Oliver J. Bosch:** conceptualization; methodology; investigation; resources; supervision; validation; project administration; funding acquisition; writing – review & editing.

## Declaration of competing interest

The authors declare no conflicts of interest.

## Notes

### Competing Interest Statement

The authors have declared no competing interest.

### Summary of Updates

The whole text has been revised to correct mistakes and further references have been included. Figures 5, 6 and 7 have been revised.

