## Supplementary for "Pup defence in lactating rats: The underlying neuropeptide signalling and their interactions in the nucleus accumbens shell" (from another version of this preprint: version 1, 2024-10-29)

### Supplementary materials

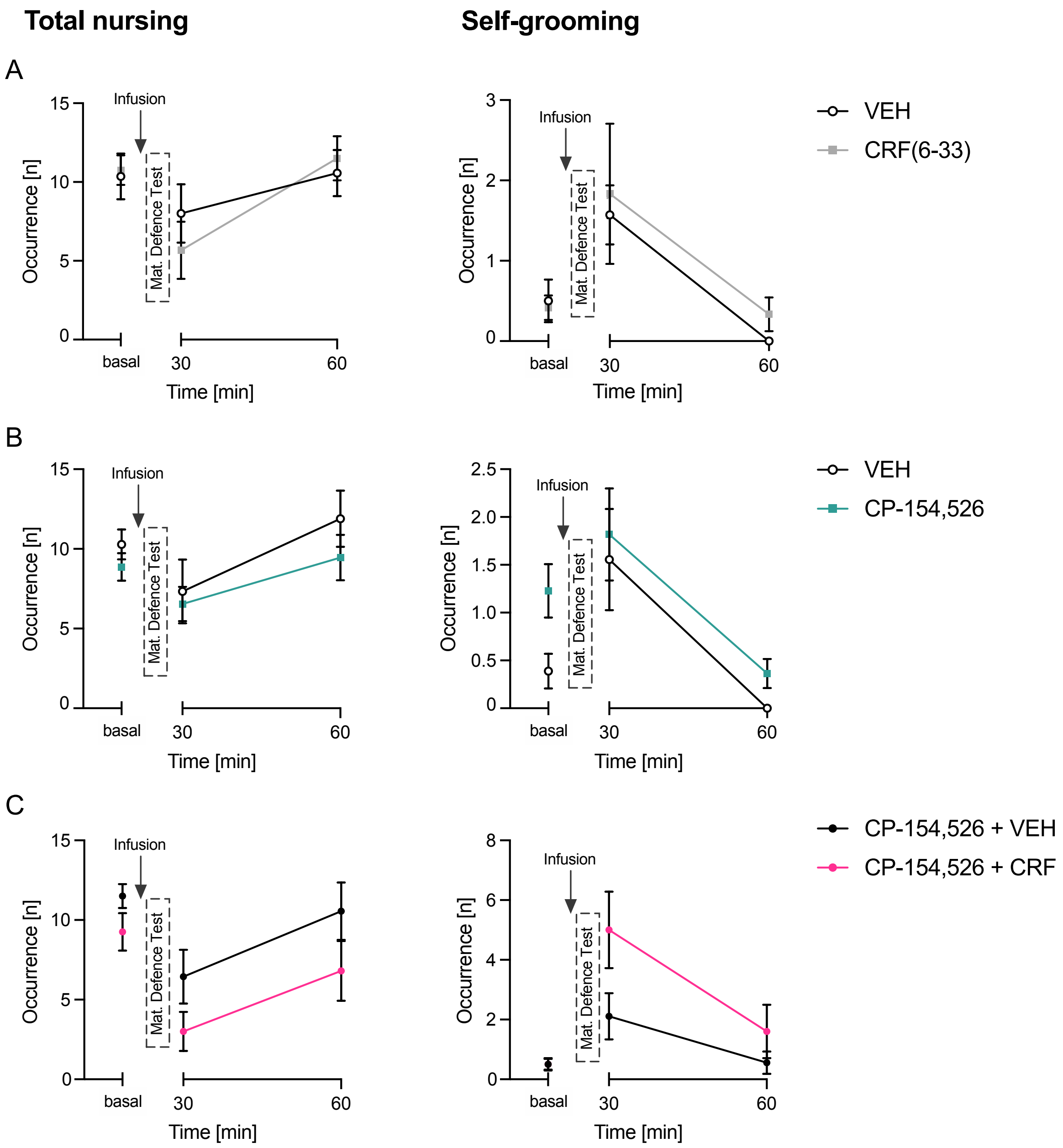


**Supplementary Figure S1. Maternal care evaluation after MDT exposure.**

Left panel shows total nursing and right panel shows self-grooming behaviour in rat mothers treated with (A) CRF(6-33) (CRF-BP inhibitor), (B) CP-154,526 (CRF-R1 antagonist), (C) CP-154,526 + CRF.

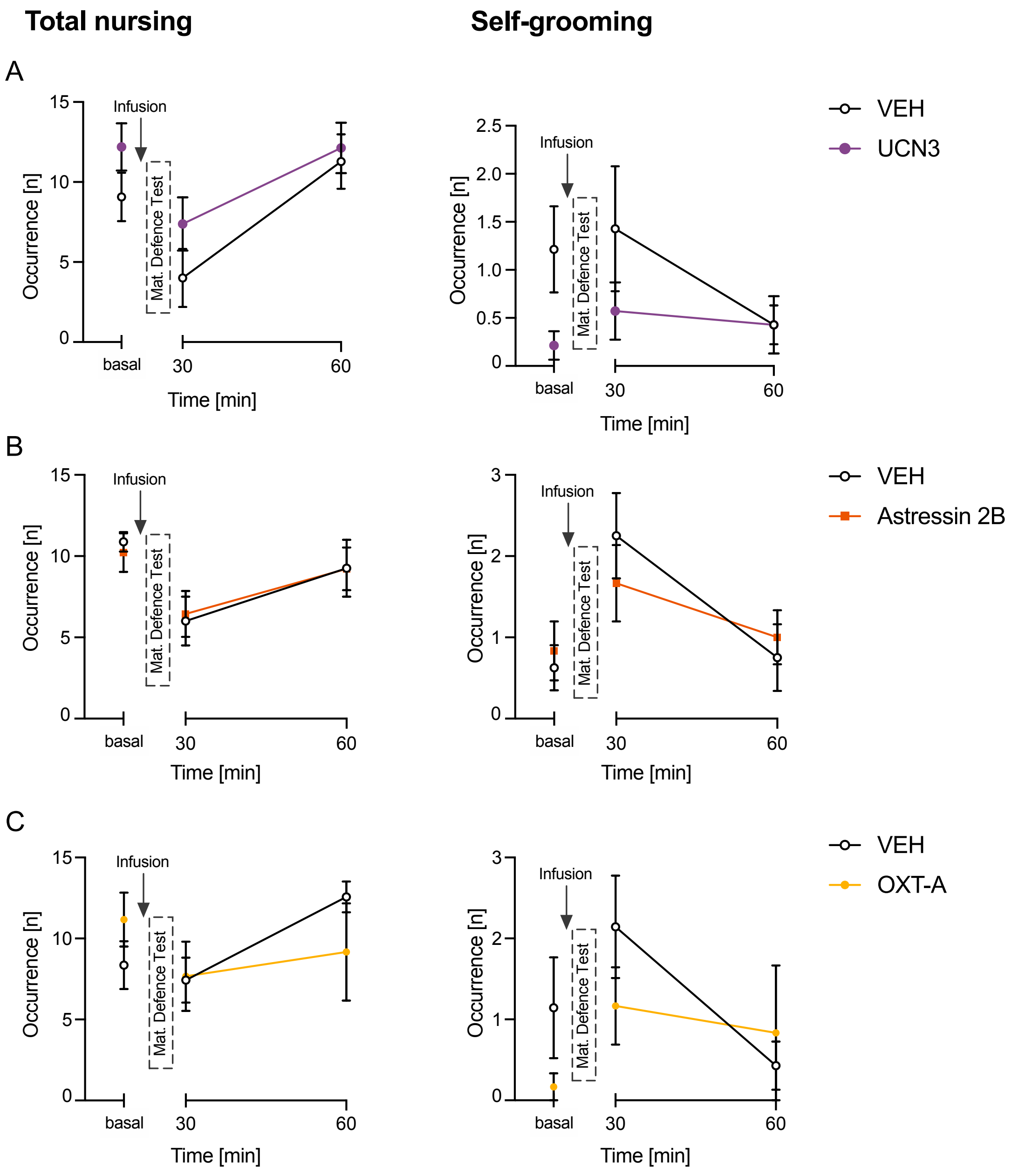


**Supplementary Figure S2. Maternal care evaluation after MDT exposure.**

Left panel shows total nursing and right panel shows self-grooming behaviour in rat mothers treated with (A) UCN3, (B) Astressin 2B (CRF-R2 antagonist), and (C) OXT-A.

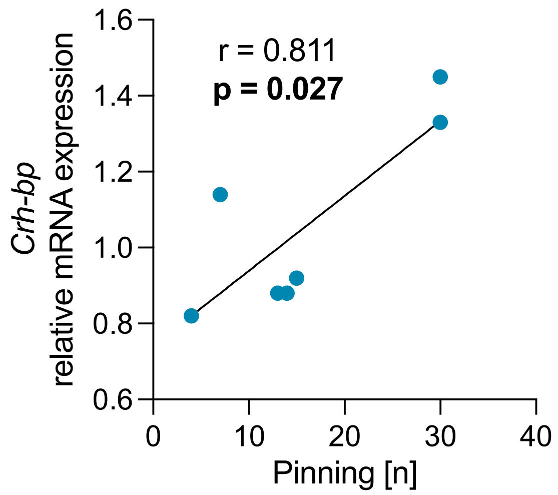


**Supplementary Figure S3. Pearson's correlation between *Crh-bp* mRNA expression levels and the number of pinning**

Relative mRNA expression levels of *Crh-bp* mRNA in the NAc showed a positive correlation with the number of pinning during the MDT in *MDT – without pups* group (Pearson's correlation). Data are presented as mean ± SEM.

**Supplementary Table S1. Statistical analysis of maternal care (i.e., nursing) after stress.** Data were analysed with 2-way repeated measured ANOVA, factors: time and treatment.

| ***Exp. Treatment*** | ***Factor Time*** | ***Factor Treatment*** | ***Factors' Interaction*** |
|---|---|---|---|
| CRF-BP inh | F[1, 17] = 5.6 | F[1, 11] = 0.005 | F[2, 22] = 0.082 |
| | **p = 0.017** | p = 0.83 | p = 0.45 |
| CRF-R1 ant | F[2, 33] = 5.7 | F[1, 18] = 1.15 | F[2, 36] = 0.269 |
| | **p =0.008** | p = 0.3 | p = 0.76 |
| CRF-R1 ant + CRF | F[2, 29] = 11.6 | F[1, 17] = 4.04 | F[2, 34] = 0.22 |
| | **p = 0.0004** | ***p = 0.061*** | p = 0.81 |
| CRF-R2 ago | F[2, 23] = 8.7 | F[1, 13] = 2.76 | F[2, 26] = 0.41 |
| | **p = 0.002** | p = 0.12 | p = 0.67 |
| CRF-R2 ant | F[2, 27] = 6.8 | F[1, 15] = 0.003 | F[2, 30] = 0.105 |
| | **p = 0.005** | p = 0.95 | p = 0.90 |
| OXT-R ant | F[1, 14] = 1.74 | F[1, 11] = 0.01 | F[2, 22] = 1.48 |
| | p = 0.21 | p = 0.94 | p = 0.25 |

**Supplementary Table S2. Statistical analysis of self-grooming after stress.** Data were analysed with 2-way repeated measured ANOVA, factors: time and treatment.

| *Exp. Treatment* | *Factor Time* | *Factor Treatment* | *Factors' Interaction* |
|---|---|---|---|
| CRF-BP inh | F[1, 12] = 9.7 | F[1, 11] = 0.22 | F[2, 22] = 0.18 |
| | **p = 0.007** | p = 0.65 | p = 0.83 |
| CRF-R1 ant | F[1, 24] = 9.8 | F[1, 18] = 3.6 | F[2, 36] = 0-4 |
| | **p = 0.002** | p = 0.074 | p = 0.67 |
| CRF-R1 ant + CRF | F[2, 28] = 14.6 | F[1, 17] = 2.54 | F[2, 34] = 2.96 |
| | **p = 0.0001** | p = 0.13 | ***p = 0.065*** |
| CRF-R2 ago | F[2, 21] = 1.47 | F[1, 12] = 2.75 | F[2, 24] = 1.32 |
| | p = 0.25 | p = 0.12 | p = 0.29 |
| CRF-R2 ant | F[2, 25] = 6.89 | F[1, 15] = 0.01 | F[2, 30] = 0.84 |
| | **p = 0.006** | p = 0.92 | p = 0.44 |
| OXT-R ant | F[2, 18] = 1.99 | F[1, 11] = 0.68 | F[2, 22] 0.92 |
| | p = 0.17 | p = 0.74 | p = 0.41 |

**Supplementary Table S3. Statistical analysis of Real-time qPCR.** Data were analysed with 2-way ANOVA, factors: test (MDT), pup presence .

| *Gene* | *Factor Pups* | *Factor Test* | *Factors' Interaction* |
|---|---|---|---|
| *Crh* | F[1, 20] = 1.0 | F[1, 20] =. 1.0 | F[1, 20] = 1.7 |
| | p = 0.32 | p = 1.65 | p = 0.20 |
| *Crh-r1* | F[1, 21] = 0.49 | F[1, 21] = 0.89 | F[1, 21] = 0.06 |
| | p = 0.49 | p = 0.36 | p = 0.80 |
| *Crh-r2* | F[1, 20] = 0.66 | F[1, 20] = 1.1 | F[1, 20] = 0.94 |
| | p = 0.43 | p = 0.3 | p = 0.34 |
| *Crh-bp* | F[1, 21] = 1.14 | F[1, 21] = 2.91 | F[1, 21] = 0.48 |
| | p = 0.30 | p = 0.1 | p = 0.49 |
| *Oxt-r* | F[1, 21] = 0.06 | F[1, 21] = 1.04 | F[1, 21] = 0.60 |
| | p = 0.81 | p = 0.32 | p = 0.44 |
